# Revealing the Distribution of Transmembrane Currents along the Dendritic Tree of a Neuron with Known Morphology from Extracellular Recordings

**DOI:** 10.1101/141069

**Authors:** Dorottya Cserpán, Domokos Meszéna, Lucia Wittner, Kinga Tóth, István Ulbert, Zoltán Somogyvári, Daniel K. Wójcik

## Abstract

Revealing the membrane current source distribution of neurons is a key step on the way to understanding neural computations, however, the experimental and theoretical tools to achieve sufficient spatiotemporal resolution for the estimation remain to be established. Here we address this problem using extracellularly recorded potentials with arbitrarily distributed electrodes in a neuron of known morphology. We use simulations of models with varying complexity to validate the proposed method and to give recommendations for experimental applications. The method is applied to in vitro data from rat hippocampus.

## 1 Introduction

Several approaches to study neurons are used in electrophysiology. Today, to monitor membrane potential, we most commonly use patch-clamp techniques [1]. Despite their unquestionable utility it is very challenging to monitor activity of a cell in more than one or two points. Extracellular recordings, on the other hand, deliver a more global picture of neural activity [2, 3]. With modern multielectrodes and microelectrode arrays one has thousands of channels at one’s disposal to monitor the brain [4–6]. However, we can no longer follow membrane potential directly but rather see spiking activity of individual cells (single-unit activity, SUA), multiple cells (multiunit activity or MUA, which is the mean firing rate of cell populations), or the mainly postsynaptic activity visible through low frequencies (so-called local field potential, LFP); see [2, 3] for discussion.

So far, the main advantages of growing throughput have been a better resolution in spike detection [7], as more cells can be identified in a single recording, improved stimulation precision [8, 9], of particular importance for retinal neuroprosthetics, and new features observed in slow fields’ profiles [10]. Recently, high density probes have been used in axon tracking [11, 12] and in studies of multisynaptic integration [13].

In the present work we ask a question, which we believe has not been posed before, although the necessary data are accessible experimentally today [13], and we propose a method to address it. Consider an acute slice where we patch a cell with a glass pipette and we drive it with intracellular current of specified time course. For simplicity, let us assume oscillatory drive, which can be subthreshold or superthreshold. Simultaneously extracellular potential is monitored with a multielectrode which will reflect the activity of the whole network, however, the contributions from the patched cell will be tied to the drive (we neglect for now possible synchronous activity of postsynaptic and other cells). Once the recordings are done, we inject a dye into the cell and reconstruct its morphology. Thus we have a set of synchronous multichannel extracellular recordings reflecting activity of a single cell whose morphology is also known, as well as its relative position to the electrode contacts. Can we use it to infer information on cell dynamics on the level of the membrane?

The traditional use of such multielectrode recordings has been to identify more and discriminate better spiking neurons [4–6, 14] or reconstruct the density of current sources (CSD) behind the recorded LFP [15–17], although more specific methods were also devised [18–20]. There were several attempts to localize cells using multielectrode recordings in different ways, taking into account the properties of electric field propagation in the tissue [14], that form the basis of CSD methods [21, 22], or other triangulation approaches [23, 24]. We are not aware of any prior attempts, however, to reconstruct current source density of individual cells using their available morphologies (although see [25]), which we propose here.

The *single cell kernel Current Source Density* method (skCSD) we introduce here is an application of the framework of the *kernel Current Source Density method* [26] to the data coming from a single cell. This is done by considering current sources located only along the cell morphology. This can be done efficiently for arbitrarily complex morphologies and arbitrary electrode configurations. In the Methods section we introduce the skCSD and explain its relations to other methods of multielectrode recordings analysis. Then, we validate this method on several ground truth datasets obtained in simulations and apply it to a proof-of-concept experimental data. Finally, we discuss practical aspects and feasibility of experimental acquisition of the required data.

## 2 Methods

### 2.1 Overview of current source density reconstruction methods

#### 2.1.1 Traditional CSD

For reader’s convenience here we briefly present the basic ideas behind the traditional and recent approaches to reconstruction of current source density (CSD analysis). For a more complete review of CSD analysis see [17], for recent reviews of the relations between neural activity, current sources and the recordings see [2, 3].

The relation between current sources in the tissue and the recording potentials is given by the Poisson equation

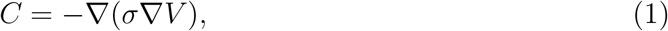

where *C* stands for CSD and *V* for the potential. While this can be studied numerically for nontrivial conductivity profiles [27], here we shall mostly assume a constant and homogeneous conductivity tensor, *σ*. In that case, the above equation simplifies to *C* = −*σ*Δ*V* and can be solved for *C* given potential in the whole space. On the other hand, given the potential in the whole space, the potential is given by

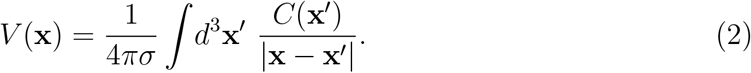

Walter Pitts observed that having recordings on a regular grid of electrodes we can estimate CSD by taking numerical second derivative of the potential [15], we call this approach *traditional CSD method.* Pitt’s idea gained popularity only after Nicholson and Freeman popularized its use for laminar recordings [28] in the cortex. In this setup, assuming the layers are infinite and homogeneous [29], the current source density at each layer can be estimated from

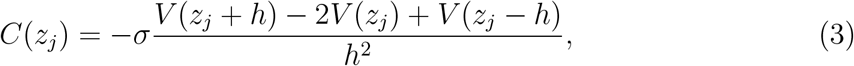

where *z_j_* is the position of the *j^th^* electrode and *h* is the inter-electrode distance.

#### 2.1.2 Inverse CSD (iCSD)

To overcome limitations of the traditional approach, such as difficulty of handling the data at the boundary and hidden assumptions about the dimensions we do not probe, Pettersen et al. proposed a model-based *inverse CSD method* [29]. Initially proposed in 1D, the method was later generalized to other dimensionalities [30, 31]. Given a set of recordings *V*_1_, …, *V_N_* at regularly placed electrodes at **x**_1_, …, **x**_*N*_ this method assumes a model of CSD parametrized with CSD values at the measurement points, 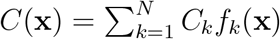, where *f_k_*(**x**) are functions taking 1 at x_*k*_, 0 at other measurement points, with the values at other points defined by the specific variant of the method, for example, spline interpolated in spline iCSD [17]. Assuming the model *C*(**x**) one computes the potential at the electrode positions obtaining a relation between the model parameters, *C_k_*, and the measured potential, *V_k_*, which can be inverted leading to an estimate of the CSD in the region of interest.

#### 2.1.3 Kernel CSD (kCSD)

The kernel Current Source Density method [26] can be considered a generalization of the inverse CSD. It is a non-parametric method which allows reconstructions from arbitrarily placed electrodes and facilitates dealing with the noise. Conceptually the method proceeds in two steps. First, one does kernel interpolation of the measured potentials. Next, one applies a “cross-kernel” to shift the interpolated potential to the CSD. In 3D, in space of homogeneous and isotropic conductivity, this amounts to applying the Laplacian to the interpolated potential, Eq. (1). To handle all cases in a general way, including data of lower dimensionality or with non-trivial conductivity, we construct the interpolating kernel and cross-kernel from a collection of basis functions. The idea is to consider current source density in the form of a linear combination of basis sources 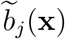, for example Gaussian,

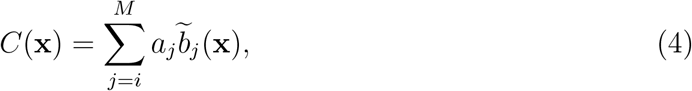

where the number of basis sources *M* ≫ *N*, the number of electrodes. Let *b_j_*(**x**) be the contribution to the extracellular potential from 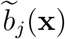, which in 3D is

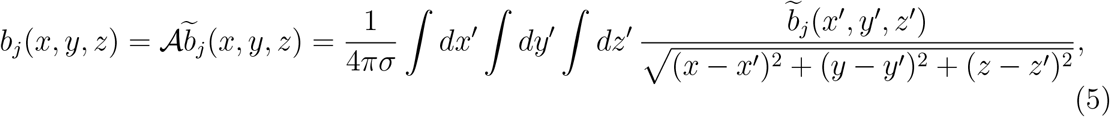

but in 1D or 2D we would need to take into account the directions we do not control in experiment (for example, along the slice thickness for a slice placed on a 2D MEA). Then, the potential will have a form

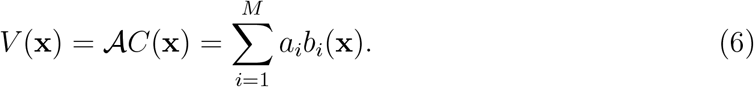

To avoid direct estimation of the coefficients *a_j_* we construct a kernel for interpolation of the potential,

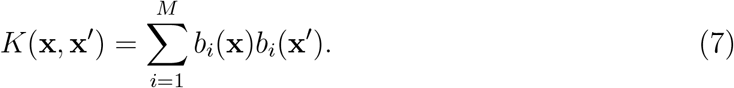

Then, any potential field *V*(**x**) span by *b_i_*(**x**) can be written as

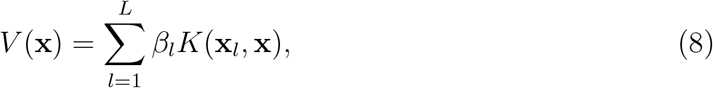

for some *L*, **x**_*l*_, and *β_l_*, but it minimizes the regularized prediction error

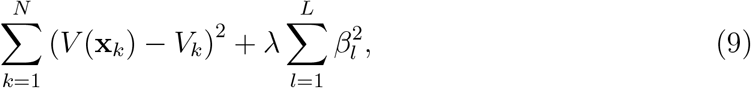

when *L* = *N*. Here, **x**_*k*_ are the positions of the electrodes, *V_k_* are the corresponding measurements, *λ* is the regularization constant. The minimizing solution is obtained for

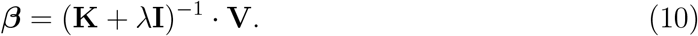

where **V** is the vector of the measurements *V_k_*, and **K**_*jk*_ = *K*(x_*j*_, x_*k*_).

To estimate CSD we introduce a cross-kernel

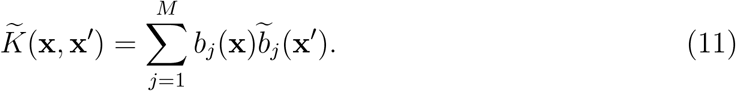

If we define

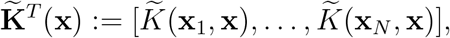

then the estimated CSD takes form of

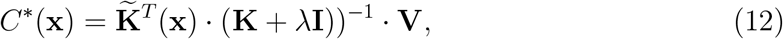

where *λ* is the regularization parameter and **I** the identity matrix; see [26] for derivation and discussion.

#### 2.1.4 Spike CSD (sCSD)

The Spike CSD [22] is the forerunner of the method presented here, as it aims to estimate the current source distribution of single neurons with unknown morphology. This requires estimation of the cell-electrode distance and a simplified model of the shape of the neuron. Separating potential patterns generated by different neurons is critical and it is obtained by clustering extracellular fingerprints of action potentials which are different for every neuron. The limitation of this model is the assumed simplified morphology of the model and low spatial resolution. Even with this simplified model it was possible to demonstrate for the first time the EC observability of backpropagating action potentials in the basal dendrites of cortical neurons, the forward propagation preceding the action potential on the dendritic tree and the signs of the Ranvier-nodes [22].

### 2.2 skCSD Method

The single cell kCSD method (skCSD), which we introduce in this work, is an application of the kCSD framework where we assume that the measured extracellular potential comes mainly from a cell of known morphology and known spatial relation to the MEA. To estimate the CSD in this case we must cover the morphology of the cell with a collection of basis functions. To do this, a one dimensional parametrization of the cell morphology is needed. This could be done independently for each branch of the neuron or globally for the whole cell at once. While the first approach might seem easier, handling of the branching point is non-trivial. Instead, we decided to fit a closed curve on the morphology, which we call the *morphology loop* (Fig. 1). This curve should cover all the segments of the cell, be as short as possible, and be aligned with the morphology. For example, in case of a ball-and-stick neuron, the curve starts at the soma, goes towards the tip of the dendrite, turns back, goes back to the soma, and closes there. One parameter s is enough to unambiguously determine a position on this line, although most points on the morphology are mapped to two *s* parameters. We also need a method to handle the branching points and guide the parametrization so that all the branches will be visited in an optimal way. This problem is a special case of the Chinese postman problem known from graph theory [32]. Given this information we can distribute the basis functions 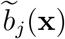 along the morphology of the cell (Fig. 1).

**Figure 1.**
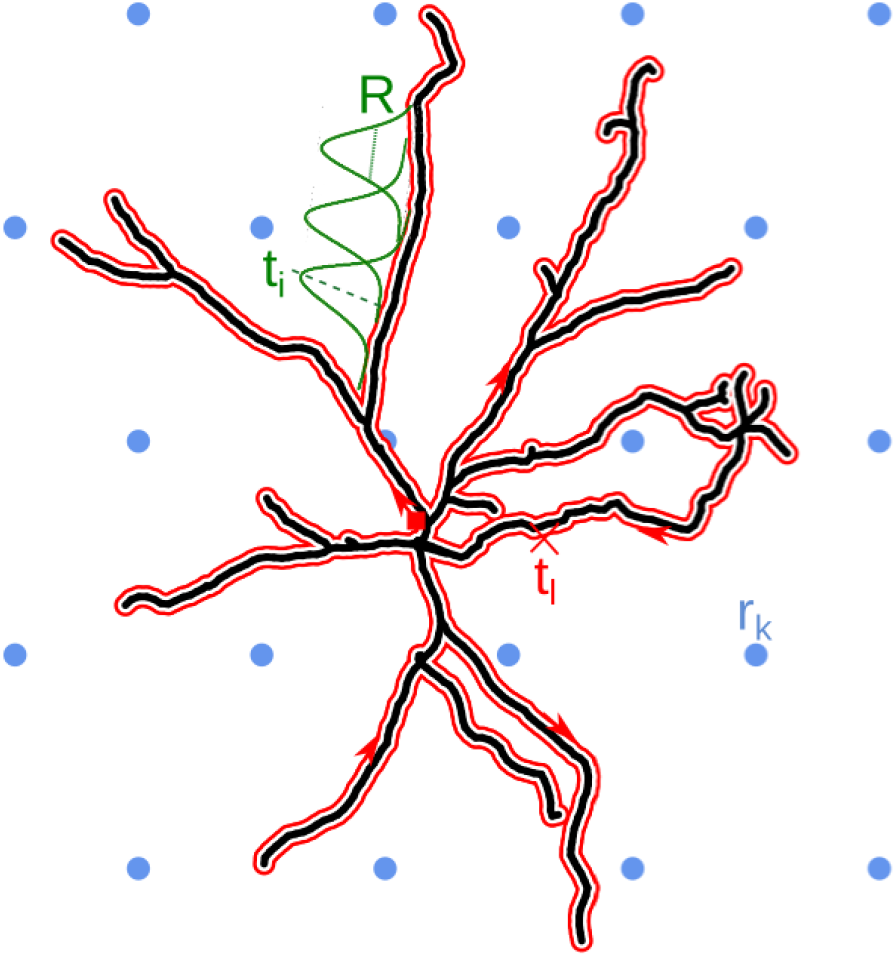
Schematic overview of the skCSD method. The black line indicates the 2-dimensional projection of the neuron on the MEA plane, the blue circles mark the location of multielectrode array (hexagonal grid, in this example), *r_k_* is the position of the *k^th^* electrode. The morphology in our method is described by a self-closing curve in three dimensions, which is indicated by red on the plot. We shall refer to this curve as the *morphology loop.* A point of the cell is visited once, if it is a terminal point of a dendrite, more than twice, if it is a branching point and twice in all the other cases. With this strategy, any point on the morphology loop uniquely identifies the physical location of the corresponding part of the cell unambiguously. To set up estimation framework we distribute 1-dimensional, overlapping Gaussian basis functions spanning the current sources. Several of these Gaussians are plotted in green, *t_i_* marks the center of the *i^th^* basis element, *R* is the width parameter.

In practice, based on the morphology information we define an ordered sequence of all the segments such that the consecutive segments are always physically connected and preference is given to those neighbors which have not been visited yet. The process is continued until all the segments are covered and the last element in the sequence connects to the first element. Note that in the sequence the final segments of the branches are present once, the branching point multiple times and the itermediate ones twice. Then we fit a spline on the coordinates of the segments following the ordered sequence resulting in a morphology loop construction. The CSD basis functions are distributed along this loop uniformly. Any point **x** ≡ (*x, y, z*) on the morphology can be parameterized with *s* ∈ [0, *l*] on the loop:

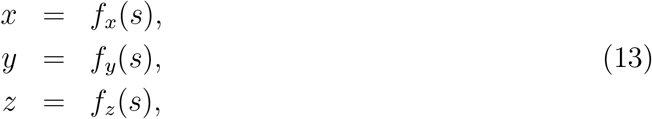

where *l* is twice the length of all the branches. Consider the following basis functions:

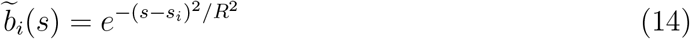

where *s_i_* is the location of the *i*-th basis function on the morphology loop, *R* its width.

The contribution to the extracellular potential from a basis source 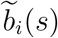 is given by

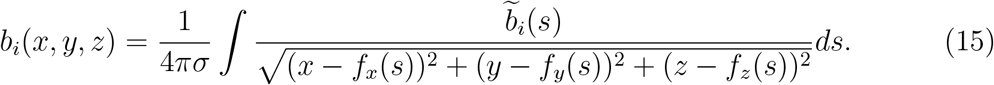

As in kCSD, for CSD of the form

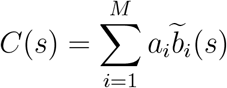

we obtain the extracellular potential as

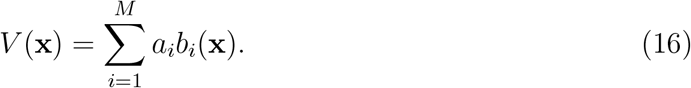

As before, for estimation of potential we use kernel interpolation. Note that in this case the basis functions in the CSD space, 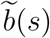, live on the morphology loop, while the basis functions in the potential space, *b_i_*(**x**), live in the physical 3D space. To determine the current source density distribution along the fitted curve we introduce the following kernel functions:

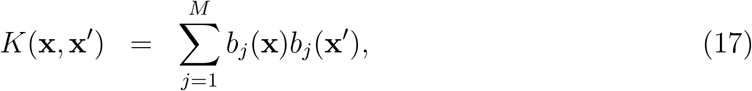

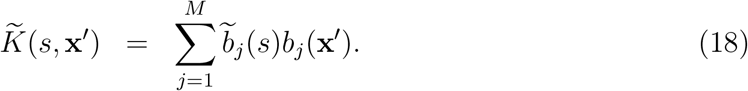

With these definitions the regularized solution for *C* on the morphology loop is given by Eq. (12):

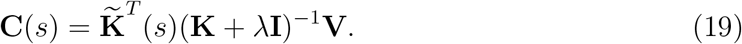

To obtain the distribution of currents at a given point in space we need to sum the currents on the loop at points which are mapped to that physical position **x**:

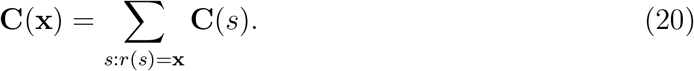

### 2.3 Construction of ground truth data

To validate the method we used simulated data which allows us to consider arbitrary cell-electrode setups and test various current patterns. The LFPy package [33] was used to simulate the extracellular potential at arbitrarily placed virtual electrodes. We assumed the `.swc` morphology description format [34] and the sections were further divided to segments. The coordinates of every segment’s ends were used to find the connections. Once the connection matrix was calculated, we used the Chinese postman algorithm to obtain the morphology loop. We calculated the potential using neuron models with various morphologies shown in Fig. 2 and different input distributions, assuming one- and two-dimensional multielectrode arrays. We used toy models to better understand and characterize the method as well as a biologically realistic neuron model to estimate performance of skCSD in an experimentally realistic scenario.

**Figure 2.**
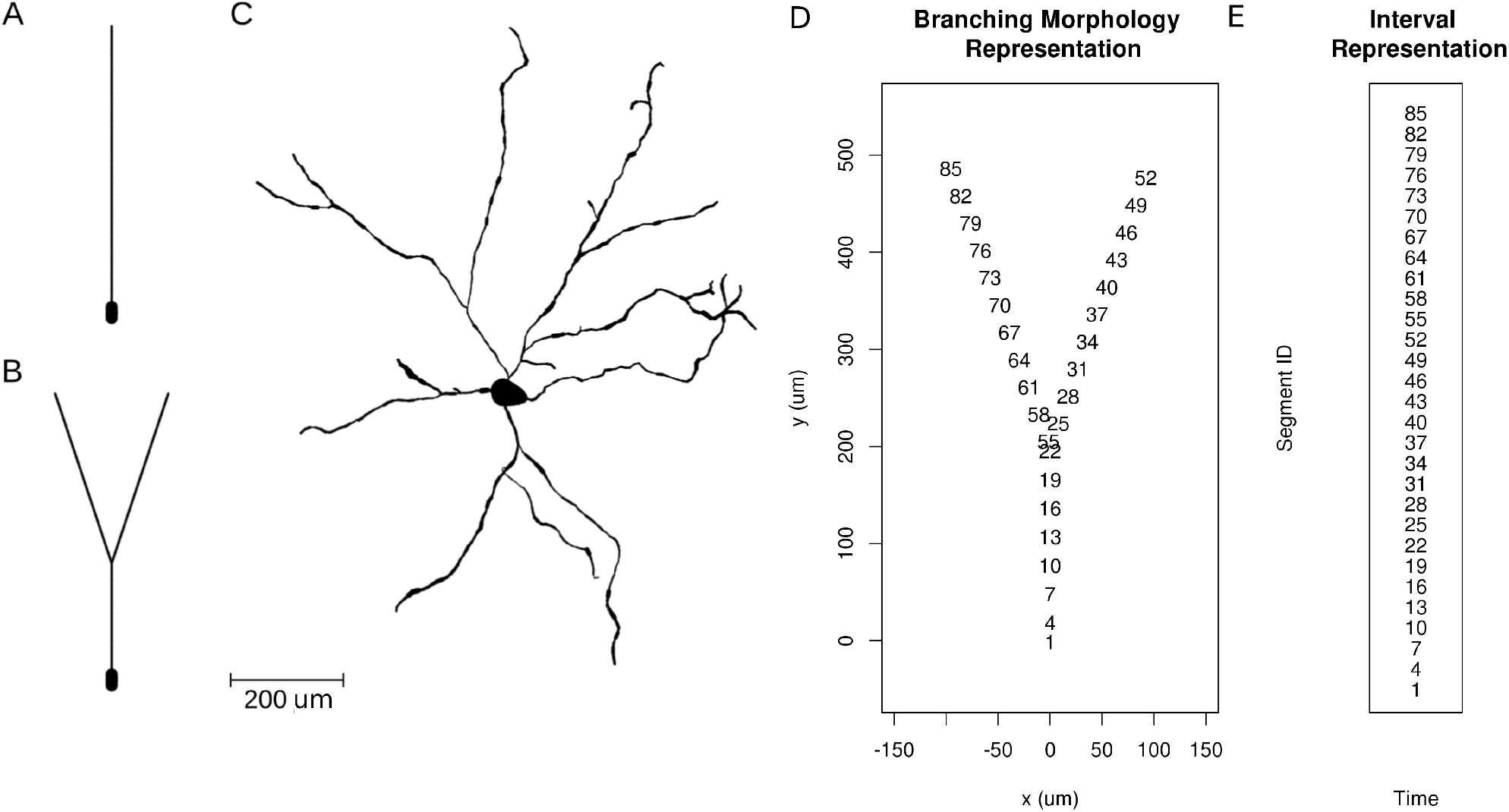
Neuron morphologies used for simulation of ground truth data. A. Ball-and-stick neuron. B. Y-shaped neuron. C. Ganglion cell.

The simplest setup we used was a ball-and-stick neuron recorded with a laminar probe. Various artificial CSD patterns and also biologically more realistic CSD distributions served as test distributions in order to quantify the spatial resolution and reconstruction errors. To generate the ground truth data we simulated a 500 *μ*m long linear cell model of 52 segments in LFPy. The diameter of the two segments representing the soma was 20 *μ*m, while the other segments were 4 *μ*m wide. 100 synaptic excitation events were distributed randomly along this morphology in order to imitate a biologically realistic scenario.

To test the effect of branching on the results, a simple Y-shaped morphology was used (Fig. 2B). The synapses were placed at segments 33 and 62 on different branches. The first was stimulated at 5, 45, 60 ms, the other at 5, 25, 60 ms after the onset of the simulation.

As a realistic example we used a mouse retinal ganglion cell morphology [35] from NeuroMorpho.Org [36]. In the simulations 608 segments were used. 100 synaptic excitation events were distributed randomly along this morphology within the first 400 ms of the simulation. The cell was also driven with an oscillatory current. In the dendrites, only passive ion channels were used.

#### Parameters of the simulations

We simulated three different model morphologies: ball-and-stick (BS), Y-shaped (Y), and a ganglion cell (Gang). The Y-shaped neuron was oncisdered in two situations, when it was parallel (Y) or orthogonal (Y-rot) to the MEA plane. The extracellular potential was computed at multiple points modeling different experimentally viable recording configurations (cell and setup). All combinations used are summarized in Table 1. The parameters describing the neuron membrane physiology are given in Table 2. The length of the simulation was 70 s in case of the ball-and-stick and Y-shaped neurons, and 850 s for the ganglion cell model.

**Table 1.** Main parameters of the simulated cells and setups.

|  | Cell Properties |  | Synapse Properties |  |  | Distribution of Electrodes |  |
| --- | --- | --- | --- | --- | --- | --- | --- |
| | Length ($\mu m$) | Number of Seg. | Location (ID of Seg.) | Number of Syn. | Synaptic Weight ($\mu S$) | Type | Number |
| **BS** | 516 | 53 | random | 100 | 0.01 | linear | 8,16, 32, 64, 128 |
| **Y** | 848 | 86 | 33, 62 | 6 | 0.04 | rectangular, random | 2x4, 4x4, 4x8, 4x16 |
| **Y-rot** | 848 | 86 | 33, 62 | 6 | 0.04 | rectangular | 8,16, 32, 64 |
| **Gang** | 5876 | 608 | random | 100 | 0.01 | hexagonal, rectangular | 128, 25, 49, 81, 441 |

**Table 2.** Biophysical parameters characterizing the simulated cell models.

| Quantity | Value | Unit |
| --- | --- | --- |
| Initial potential | -65 | $mV$ |
| Axial Resistance | 123 | $\Omega cm$ |
| Membrane resistivity | 30000 | $\Omega cm^2$ |
| Membrane capacitance | 1 | $\mu F/cm^2$ |
| Passive mechanism reversal potential | -65 | $mV$ |

#### Parameters of synapses

In most simulations we modeled synaptic activity. We used synapses with discontinuous change in conductance at an event followed by an exponential decay with time constant *τ* (ExpSyn model as implemented in the NEURON simulator). When simulating the Y-shaped neuron we placed two synapses with the following parameters: reversal potential: 0 mV, synaptic time constant: 2 ms, synaptic weight: 0.04 *μ*S. The synapses were placed at segments 33 and 62 (See Fig. 2 and 5). When simulating the other models (ball-and-stick and ganglion cell) we used the same type of synapse, however, the synaptic weights were a quarter of the above (0.01 *μ*S) since they were more numerous (Table 1).

### 2.4 Measuring the Quality of Reconstruction

To validate the skCSD method we need to consider two situations. When we know the ground truth — the actual distribution of sources which generated the measured potentials — we can compare the reconstruction with it. This is available directly only in simulations. In that case we can measure the prediction error between the reconstruction and the original. However, the skCSD method by its nature gives smooth results. This is a consequence of kernel interpolation of the potential which occurs in the first step of the method. The same phenomenon occurs in regular CSD estimation [17]. Thus, we can never recover the original CSD distribution but only a coarse-grained approximation. This is not a significant problem as the coarse-grained CSD should have equivalent physiological consequence. However, to compare the reconstructed density with the ground-truth, which is typically very irregular in consequence of multiple synaptic activations, we always smoothed the ground truth CSD with a Gaussian kernel. The width of the kernel was 15 *μm* for ball-and-stick model, while for the Y-shaped and ganglion cell models we used 30 *μm*.

Thus, whenever ground truth was known, we computed L1 norm of the difference between the reconstruction *C*\* and smoothed ground truth *C* normalized by the L1 norm of *C*:

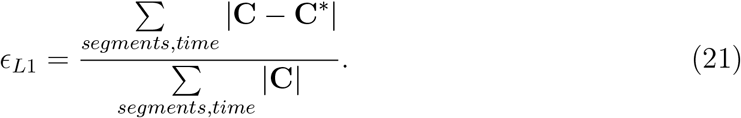

When analyzing experimental data we only have access to the noisy measurements and cannot apply the above strategy directly. Thus we consider two strategies. One is to use cross-validation error (CV). In leave-one-out cross-validation [26] we estimate CSD from all the measurements but one and compare estimated prediction with actual measurement on the removed electrode. Repeating this procedure for all the electrodes gives us a measure of prediction quality for a given set of parameters for this specific dataset. Scanning over some parameter range we identify optimal parameters as those giving minimum error. They are further used to analyze the complete data. The advantage of using cross-validation error is that it does not require the knowledge of the ground truth current source density distribution and can still provide an estimation about the performance of the skCSD method. As this algorithm is quadratic in the number of electrodes, for large arrays one might prefer to use the leave-p-out cross-validation instead. When we test how the quality of the reconstruction changes with the number of electrodes we use CV error normalized by the number of electrodes which can then be compared between different setups.

The other strategy we use and recommend in the experimental context, when we know the cell morphology and its geometric relation to the setup, as well as the measurements, is model-based analysis. The idea is to simulate different current source distributions, either placing specific distribution by hand or by modeling activity of the cell assuming passive membrane and random or specific synaptic activations, both of which are relatively inexpensive both in computational time and coding complexity. This reduces the problem to the modeling case. We can use thus generated data (CSD and potentials) scanning for optimal reconstruction parameters to be used in analysis of actual experimental data from the setup.

To handle the effects of noise one should study its properties on electrodes, e.g., assuming white measurement noise identify its variance, then tune the regularization parameter λ on simulated sets with comparable simulated noise added.

### 2.5 Parameter selection

To apply the skCSD method we need to decide upon the number of basis functions, set their width (*R*), and choose the regularization parameter *λ*. In this work the number of basis function was set to 512 for all cases, which is at least twice the number of electrodes used. This is usually not a limitation, the more the better. For the basis width (Eq. (14)) we took the following values: 8, 16, 32, 64, 128 *μm*. Selection of the regularization parameter is not trivial [26, 37]. Here, we tested the effect of the regularization parameter taking values of 0.00001, 0.0001, 0.001, 0.01, 0.1 The optimal parameters were identified by the lowest value of reconstruction error.

### 2.6 Visual representation of CSD on the morphology

To visualize the distribution of current sources and other quantities along a neuron morphology we use two representations of the cell:

1. Interval: we stack all the compartments consecutively along the y-axis so that the part of the dendrite stemming from the soma is shown first, followed by one branch, followed by the other. The order of the branches in the stack is taken from the morphology loop to make these representations consistent. The *x*-axis either shows different time instants of the simulations or various distribution patterns.
2. Branching morphology: in this case a two-dimensional projection of the cell is shown which is colored according to the amplitudes of the membrane current source densities at a time instant. To visually enhance the current events, gray circles proportional to the amplitude of CSD at a point are placed centered at the point to facilitate comprehension.

### 2.7 Experimental methods

#### In vitro experiment

One male Wistar rat (300*g*) was used for the slice preparation procedure. The in vitro experiment was performed according to the EC Council Directive of November 24, 1986 (86/89/EEC) and all procedures were reviewed and approved by the local ethical committee and the Hungarian Central Government Office (license number: PEI/001/695-9/2015). The animal was anesthetized with isoflurane (0.2 ml/100g). Horizontal hippocampal slices of 500 μm thickness were cut with a vibratome (VT1200s; Leica, Nussloch, Germany). We followed our experimental procedures developed for human in vitro recordings [38], adapted to rodent tissue. Briefly, slices were transferred to a dual superfusion chamber perfused with artificial cerebrospinal fluid. Intracellular patch-clamp recordings, cell filling, visualization and three-dimensional reconstruction of the filled cell was performed as described in [38]. For the extracellular local field potential recordings, we used a 16-channel linear multielectrode (A16x1-2mm-50-177-A16, Neuronexus Technologies, Ann Arbor, MI, USA), with an INTAN RHD2000 FPGA-based acquisition system (InTan Technologies, Los Angeles, CA, USA). The system was connected to a laptop via USB 2.0. Wideband signals (0.1—7500 Hz) were recorded with a sampling frequency of 20 kHz and with 16-bit resolution. The recorded neuron was held by a constant −40 nA current injection.

#### Data preprocessing

154 spikes were detected on the 180s long intra-cellular recording by 0 *mV* upward threshold crossing. A ±5 s wide time windows were cut around the moments of each spikes on each channels of the extra-cellular (EC) potential recordings and averaged, to access the fine details of the EC spatio-temporal potential pattern which accompanied the firing of the recorded neuron on all channels. Two channels were broken (2, 5), however, as the skCSD method allows retrieving CSD maps from arbitrarily distributed contacts, this has not prevented the analysis; the broken channels were excluded from further consideration. The averaged spatio-temporal potential maps were high-pass filtered by subtracting a moving window average with 100 ms width. This filtering, together with the spike triggered averaging procedure, ensured that the resulted EC potential map contains only the contribution from the actually recorded cell. The price we paid was filtering out EC signals of the spontaneous repetitive sharp-wave like activity of the slice which was correlated by the firing of the recorded neuron and thus the presumptive synaptic inputs of the recorded neuron as well. An additional temporal smoothing by a moving average with 0.15 ms window was used to reduce the effect of noise.

## 3 Results

In this section we study the properties of the skCSD reconstruction for three representative morphologies of increasing complexity and for different setups. First, for a ball-and-stick neuron, we study the general quality of reconstruction of fine detail by considering oscillating CSD distributions of increasing spatial frequency which form the Fourier basis. Since the oscillating sources are not a natural representation for branching morphologies, there we show examples of reconstructions for random or specific activation, typically synaptic, which might arise in experimental context. To build intuition on how the Fourier space representation translates into a specific distribution we consider reconstruction of sources for random synaptic activation of the ball-and-stick cell.

Then, for a neuron with a single branching point (Y-shaped morphology), we check if skCSD can differentiate between synaptic activations close to the branching point located on different branches. We also investigate the effects of random electrode placement on skCSD reconstruction. Finally, we investigate the possibility of skCSD reconstruction on a realistic model of a ganglion cell placed on a MEA as well as the sensitivity to noise of the method.

After establishing and validating the method on these fully controlled model data, to show experimental viability of the proposed method, the spike-triggered average current source density distribution is reconstructed for a pyramidal cell from experimental data.

### 3.1 Ball-and-stick neuron

Here we consider the simplest neuron morphology, so-called ball-and-stick model, which stands for the soma and a single dendrite. A virtual linear electrode was placed in parallel to the cell 50 *μm* away, the electrodes were distributed evenly along the electrode extending for 600 *μm*.

#### Increasing the density and number of electrodes improves spatial resolution of the method

To study the spatial resolution of the skCSD method we consider ground truth membrane current source density distributions in the form of waves with increasing spatial frequencies

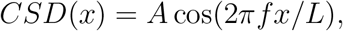

where *A* = 0.15 *nA*/*μm* is the amplitude, *f* ∈ {0.5, 1, 1.5, …, 12.5} is the spatial frequency, *x* is the position along the cell, *L* is the length of the cell. Then, we compute the generated extracellular potential at the electrode locations. The laminar shank consisting of 8, 16 and 128 electrodes was placed 50 *μm* from the cell in parallel to the dendrite. Finite sampling of the extracellular space sets a limit on the spatial resolution of the method. Increasing the density of electrodes within the studied region leads to higher spatial precision. As we can see in Fig. 3, with 128 electrodes it is possible to reconstruct higher frequency distributions as compared to 8 electrodes. This is reminiscent of the Nyquist theorem, except here we measure the potential and reconstruct current sources, while Nyquist applies to sampling and reconstruction of the same quantity. What we observe is quite intuitive and typically observed in the discrete inverse methods [37]. Note that once we move to complex morphologies and random rather than regular electrode placement, the intuition we build here, that denser probing gives better spatial resolution, would still hold, even if the relation with the Nyquist theorem would be less apparent.

**Figure 3.**
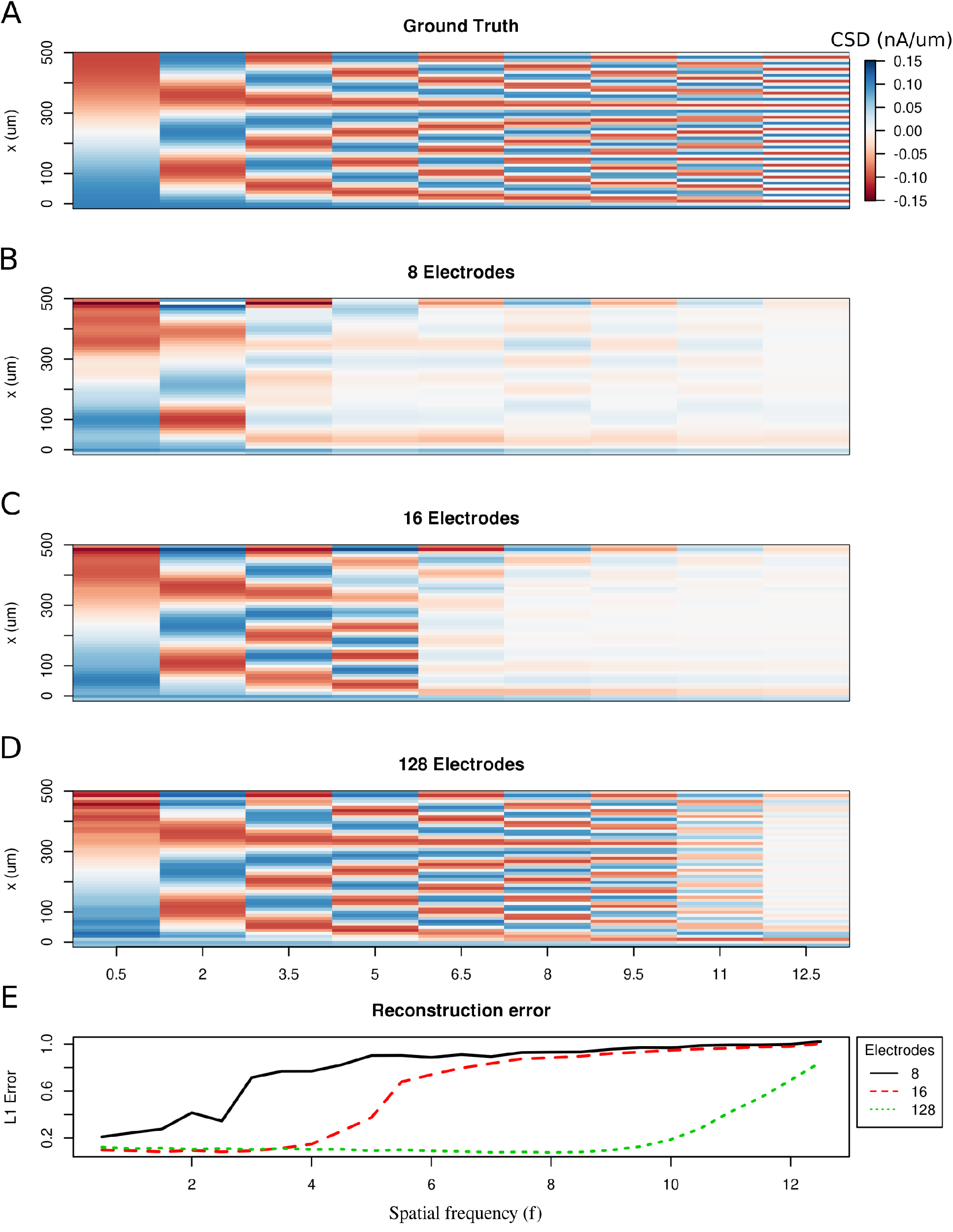
Limitations of the spatial resolution of the skCSD method in a simple ball-and-stick and laminar electrode setup. **A**. The ground truth membrane current source density distribution was constructed from cosine waves of increasing spatial frequency (x-axis) along the cell mophology (y-axis), which is shown in the interval representation. **B–D**. skCSD reconstruction from 8, 16 and 128 electrodes. **E**. The L1 Error of the skCSD reconstruction for 8 (black), 16 (red) and 128 (green) electrodes for CSD patterns of increasing frequency.

#### Reconstruction of random synaptic activations

In the same setup as before we place 100 synapses along the dendrite and stimulate them randomly in time. We simulate 70 ms of recordings from this synaptically activated cell. The stimulation is sufficiently strong to evoke spiking, see the Appendix for details. The spiking is indicated by strong red spots in the lowest first two segments in Fig. 4, which correspond to the soma. As we can see, the reconstructed current source density distribution reflects the ground-truth, and the precision of reconstruction improves with increasing the number of electrodes, which is reflected in the reduction of cross-validation error. Notice how the reconstructed synaptic activity gets more precise with increasing the density of probing the potential.

**Figure 4.**
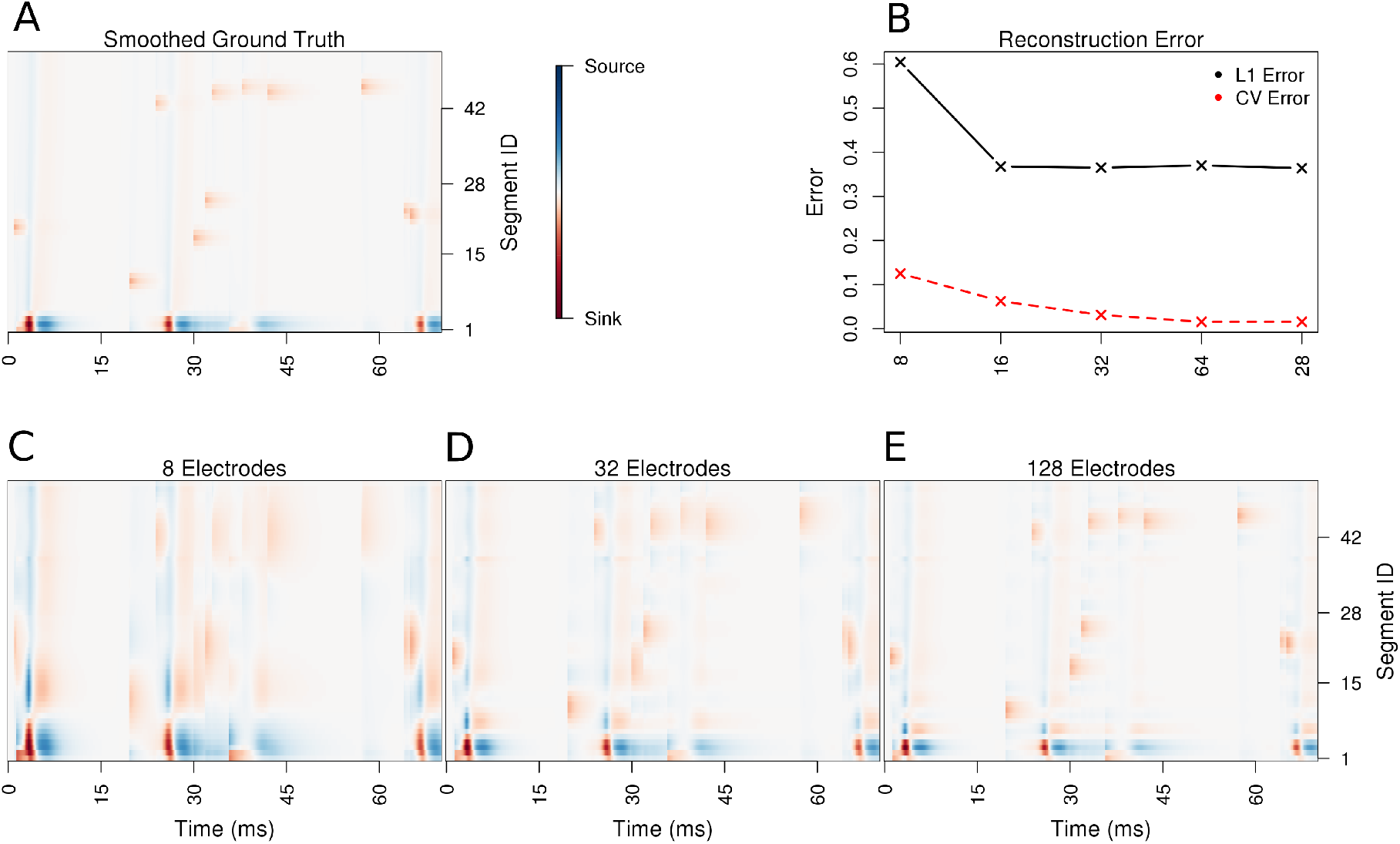
Performance of the skCSD method for a ball-and-stick neuron with random synaptic stimulation for recordings with a laminar probe placed 50 *μm* away from the cell. **A**. The ground truth spatio-temporal membrane current density in time (x-axis) along the cell in the interval representation (y-axis). The lowest segment is the soma, where the visible high amplitude of potential is a consequence of spiking. To make the much less pronounced synaptic activity on the dendritic part visible, nonlinear color map was used. Panel **B** shows the lowest values of cross-validation and L1 error for the before-mentioned setups. Panels **C–E** present the best skCSD reconstruction in case of recording with 8, 32, and 128 electrodes. One can see how increasing the number and density of probes in the region improves the reconstruction quality until a certain level. CV error was used here to select the parameters leading to the best reconstructions.

### 3.2 Simple branching morphology

Let us now study the effect of branching and breaking of rotational symmetry of the cell on the skCSD method. We consider here a simple *Y* -shaped model neuron with one branching point (Fig. 2 B). We place two synapses, one on each branch (at segments 33 and 62, close to the branching point, see Fig. 2 D and Fig. 5 C). We consider both simultaneous and independent activation of these synapses, specifically, the first synapse was activated at 5, 45, 60 ms of the 70 ms long simulation, while the other was stimulate at 5, 25, 60 ms from the stimulation onset. Our goal here is to find out if we can separate the synaptic inputs located on two different branches, what happens at the branching point, how the arrangement of the electrodes-cell setup affects the reconstruction, and if our method provides more detail about the current distribution on the cell than what is accessible from the interpolated potential and the CSD reconstructed with kCSD under the assumption of smooth distribution of sources in space.

**Figure 5.**
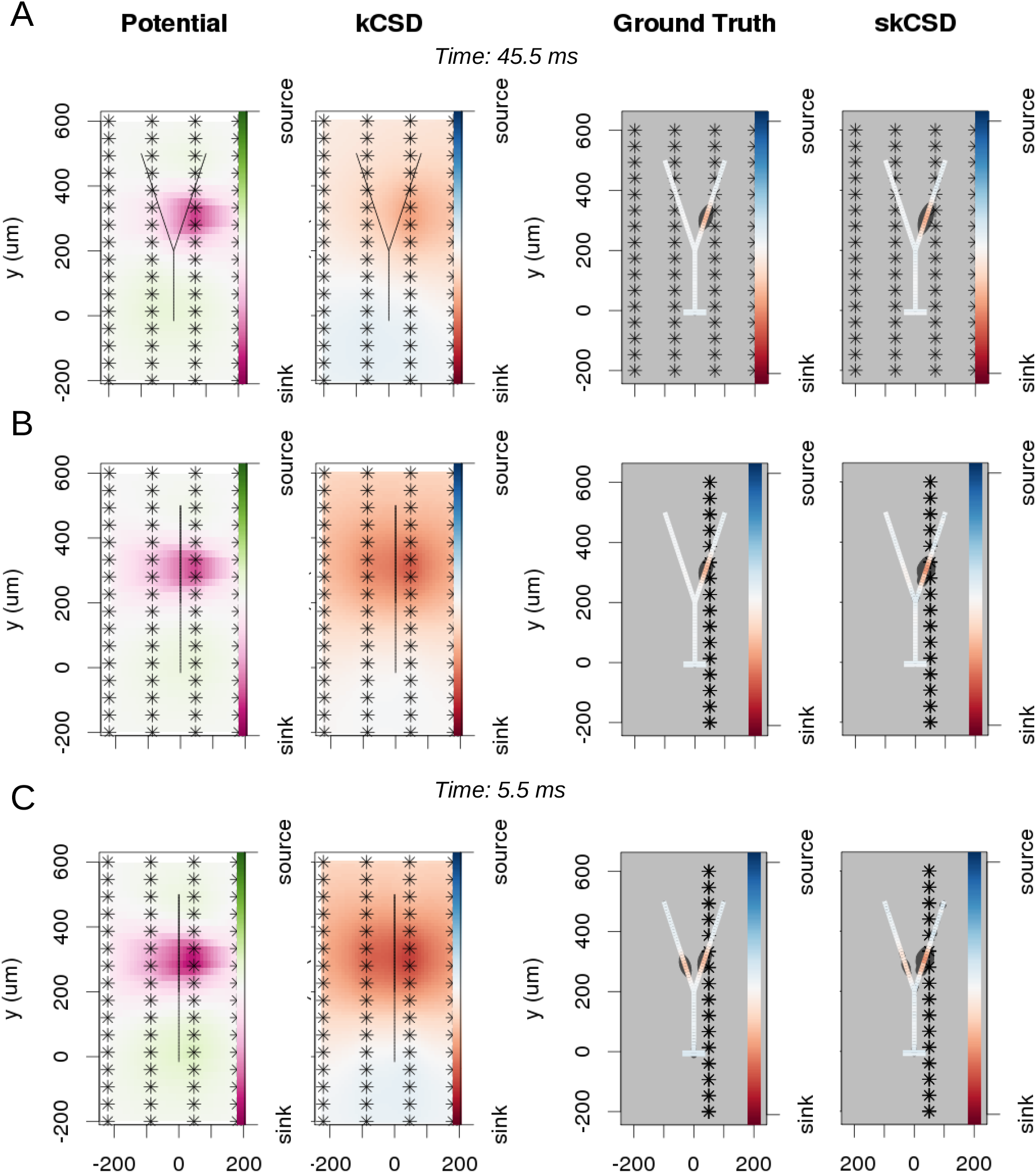
Reconstruction of synaptic inputs on a Y-shaped neuron with a regular rectangular 4×16 electrode grid. Each panel (**A–C**) shows the spline-interpolated extracellular potential (V), followed by standard kCSD reconstruction, both at the plane of the 4×16 electrodes’ grid used for simulated measurement. Then, the ground truth and skCSD reconstruction are shown in the branching morphology representation in the plane containing the cell morphology. Each figure shows superimposed morphology of the cell. Note that in panel **A** the grid is parallel to the cell, while in panels **B–C** it is perpendicular. The dark gray shapes are guides for the eye and are sums of circles placed along the morphology with radius proportional to the amplitude of the sources at the center of the circle. **A**. Shows results for a synaptic input depolarizing one branch. **B**. Shows the same current distribution as in the previous setup, but the grid is rotated by 90 degrees. **C A** synaptic input is added to the other branch. Observe that in all three cases the interpolated potential and the standard CSD reconstruction, which can be drawn only in the plane of the electrodes’ grid, do not differ significantly, hence they cannot distinguish between these three situations. On the other hand, skCSD method is able to identify correctly both synaptic inputs.

#### Differentiation of synaptic inputs located on different branches

To investigate the differentiation power of the proposed approach we consider two placements of the cell with respect to the electrode grid. One, in which the cell is placed in parallel to the plane of electrodes 50*μm* above (Fig. 5 A), and the other, where the cell is perpendicular to the grid, with the grid 50*μm* away from the dendritic shaft stemming from the soma, (Fig. 5 B, C). In Fig. 5, each panel (**A–C**) shows the spline-interpolated extracellular potential (V), followed by standard kCSD reconstruction, both at the plane of the 4x16 electrodes’ grid used for simulated measurement. Then, the ground truth and skCSD reconstruction are shown in the branching morphology representation in the plane containing the cell morphology. Each figure shows superimposed morphology of the cell. The dark gray shapes are guides for the eye and are sums of circles placed along the morphology with radius proportional to the amplitude of the sources located at the center of the circle. Panel **A** shows results for a synaptic input depolarizing one branch. Panel **B** shows the same current distribution as in the previous setup, but the cell is rotated by 90 degrees with respect to the grid. In panel **C** a synaptic input is added to the other branch. Observe that in all three cases the interpolated potential and the standard CSD reconstruction, which can be drawn only in the plane of the electrodes’ grid, do not differ significantly, hence they cannot distinguish between these three situations. On the other hand, skCSD method is able to identify correctly the synaptic inputs in all three cases.

#### The effect of electrodes placement on skCSD reconstruction for Y-shaped cell

In Fig. 6 we show how the number and specific distribution of the electrodes affect the quality of the reconstruction in the case of simultaneous stimulation. Panel 6. **A** shows the ground truth data, that is the actual distribution of the transmembrane current sources, along the morphology. To visualize it simply, we used the interval representation, the soma is shown first, followed by one branch, followed by the other. Fig. 6.**B** shows the reconstruction results for regularly arranged 8 (4×2), 16 (4×4), 32 (4×8), and 64 (4×16) electrodes. In Fig 6.**C** we show reconstructions for five different random placements of the same number of electrodes as for the regular case. As expected, the skCSD method is able to recover the synaptic activations and the reconstruction resolution increases with the number of electrodes. Note that in certain cases the random distribution is more efficient than the regular grid, which is probably due to more fortunate samplings of the area covered by the morphology.

**Figure 6.**
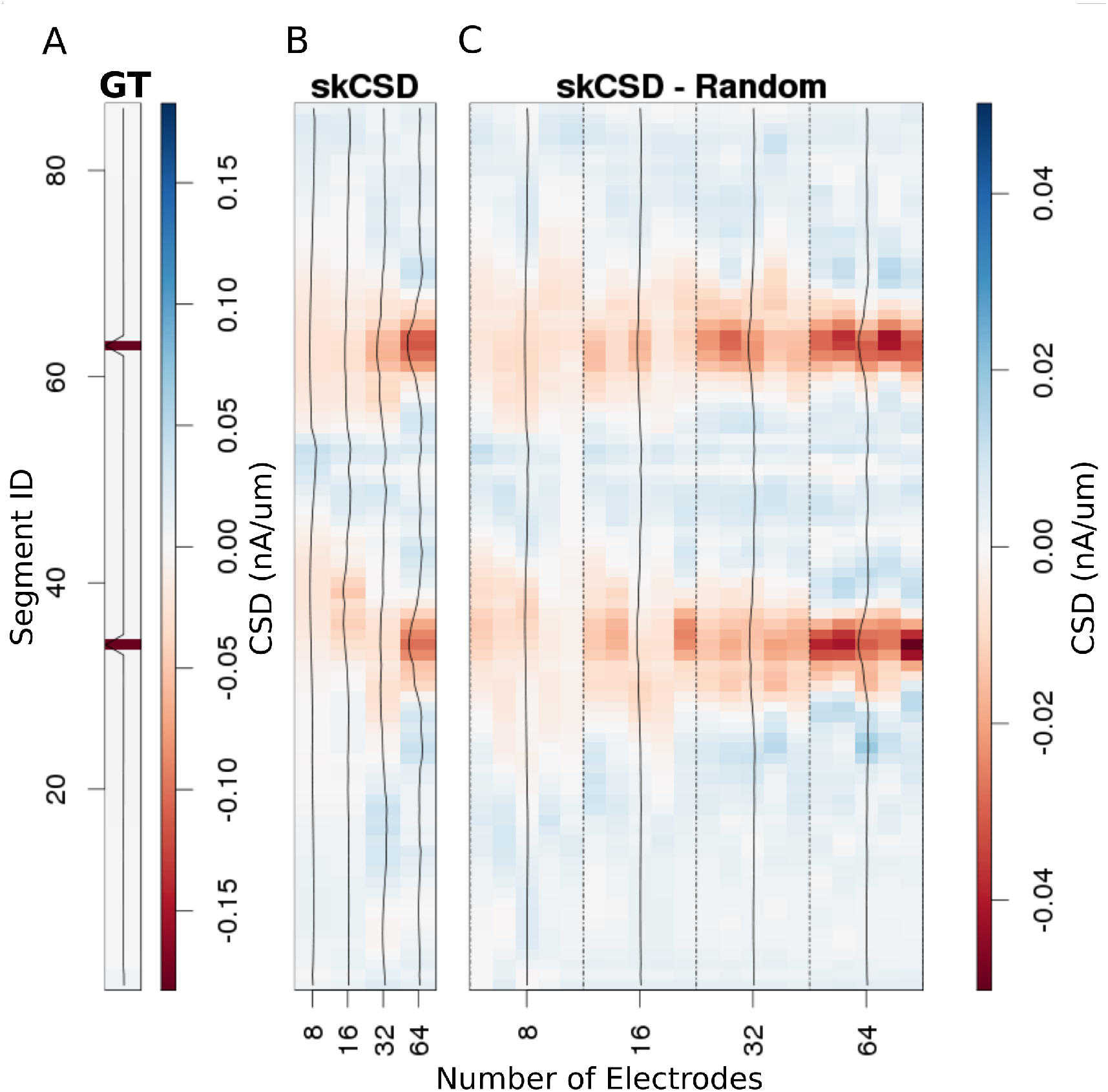
Reconstruction of synaptic inputs placed on different branches of the Y-shaped neuron for electrodes arranged regularly and randomly within the same area. We use the interval representation for visualization. The numbers on horizontal axis enumerate different electrode setups. The black profiles show the averaged membrane current as reconstructed in a given case; for random electrode distribution these are averages over five different realizations. **A** Ground truth membrane currents, the strong red indicates the synaptic inputs. **B** Reconstruction results for 8 (4×2), 16 (4×4), 32 (4×8), and 64 (4×16) electrodes arranged regularly. The skCSD reconstruction improves with the number of electrodes as the color representation and the black profiles indicate. **C** When distributing the same numbers of electrodes on the same plane as in the previous case, the quality of the average skCSD reconstruction, as indicated by the black profiles, is similar.

### 3.3 Reconstruction of current distribution on complex morphology

In this section we consider the performance of skCSD method in case of complicated, biologically realistic scenario. To reach good spatial resolution allowing detailed study of a cell with substantial extent, densely packed electrode arrays are required. In the present reconstruction we assumed a hexagonal grid arrangement with 17.5 *μm* interelectrode distance inspired by recent experiments on reconstructing axonal action potential propagation [11, 25]. We assumed the grid consisting of 936 contacts from which we used 128 electrodes for reconstruction to be consistent with the hardware of [11, 25].

In the simulation we assumed an experimentally plausible scenario, where oscillatory current was injected to the soma of a neuron in a slice with other inputs impinging through a 100 excitatory synapses distributed on the dendritic tree. The simulated data consisted of two parts. During the first 400 ms the cell was stimulated by the injected current as well as through the synapses. The amplitude of the injected current was 3.6 nA, the frequency of the current drive was around 6.5 Hz. During the second 400 ms the cell was stimulated only with the current. Fig. 7 shows an example of the skCSD reconstruction at a time selected right after a spike was elicited by the cell. As we can see, neither the standard CSD recontruction assuming smooth current distribution in space, nor the interpolated potential, give justice to the actuall current distribution. At the same time, the skCSD reconstruction is quite a faithful reproduction of the ground truth. A movie comparing the ground truth with kCSD, interpolated potential, and skCSD reconstruction, in time, is provided as a supplementary material (S1 Video).

**Figure 7.**
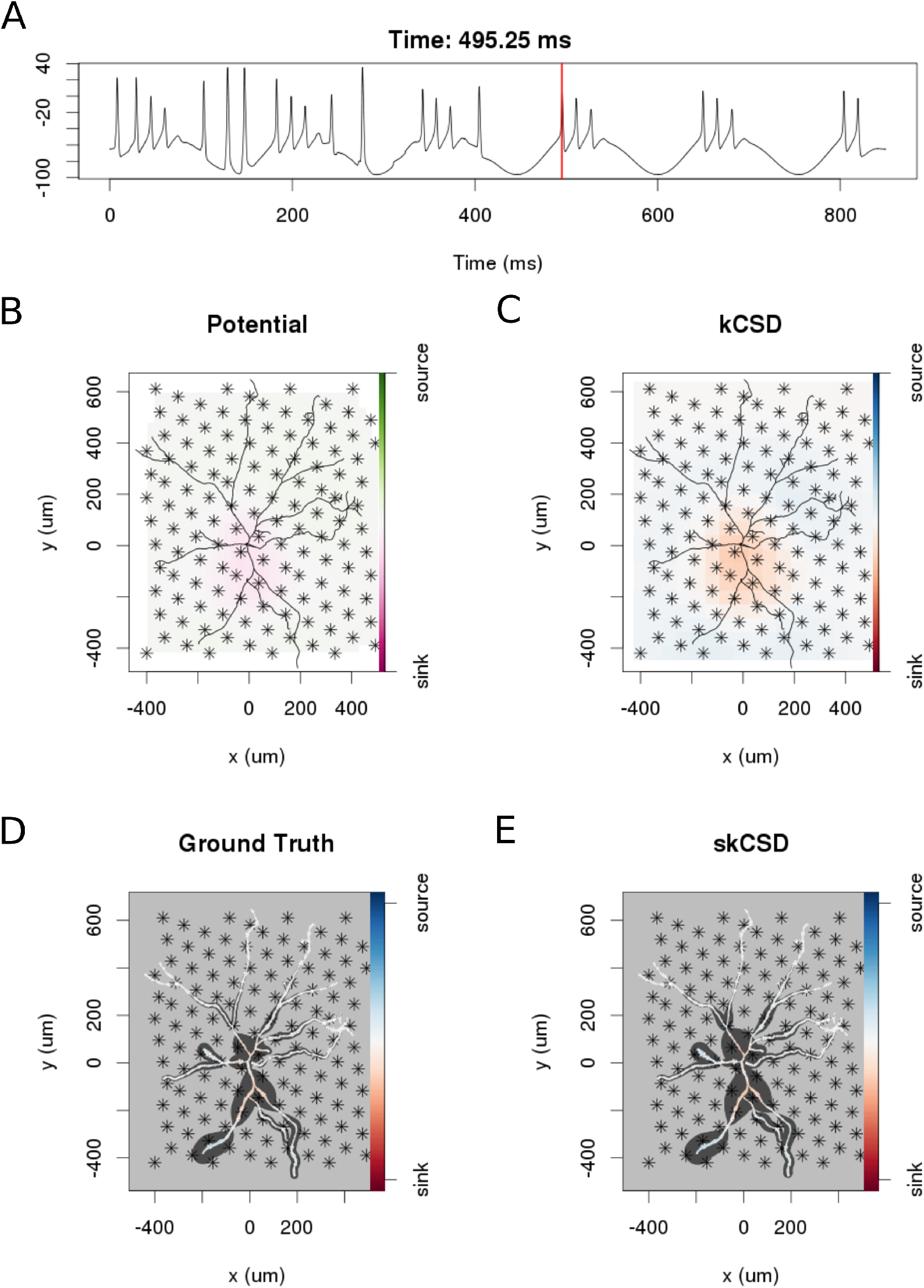
skCSD reconstruction of dendritic backpropagation patterns for a retinal ganglion cell model driven with oscillatory current. **A** Somatic membrane potential during the simulation. The red line marks the time instant for which the remaining plots were made. **B** Extracellular potential interpolated between the simulated measurements computed at the electrodes, which are marked with asterisks. **C** kCSD reconstruction computed from the simulated measurements of the potential. **D** Spatial smoothing with a Gaussian kernel was applied to the ground truth membrane current to facilitate comparison with the skCSD reconstruction with the same spatial resolution level. **E** skCSD reconstruction computed from the simulated measurements of the potential.

### 3.4 Dependence of reconstruction on noise level

So far we have assumed that the data are noise-free which is never true in an experiment. Both the measurement device and the neural tissue are potential sources of distorted data. To investigate how the performance of the method is influenced by noise, we added Gaussian white noise of differing amplitudes to the simulated extracellular recordings of Y-shaped cell described in Section 3.2. Fig. 8. A shows the smoothed ground truth we used. The Y-shaped neuron is placed on top of a MEA with a regular grid of 4×8 electrodes marked by asterisks. Fig. 8.B shows the noise-free reconstruction. Panel C–F of the figure show the reconstruction results for increasing measurement noise with signal to noise ratio, SNR= 16, 4, 1. The signal-to-noise ratio (SNR) here is the standard deviation of the simulated extracellular potentials normalized with the std of the added noise. The degradation of reconstruction visible in this figures is summarized in Fig. 8.C. As we can see in the reconstruction plots (Fig. 8.D–F), the increasing noise actually does not seem to significantly alter the obtained reconstructions so the regularization is providing adequate correction, except for the noise on the order of signal (Fig. 8.F).

**Figure 8.**
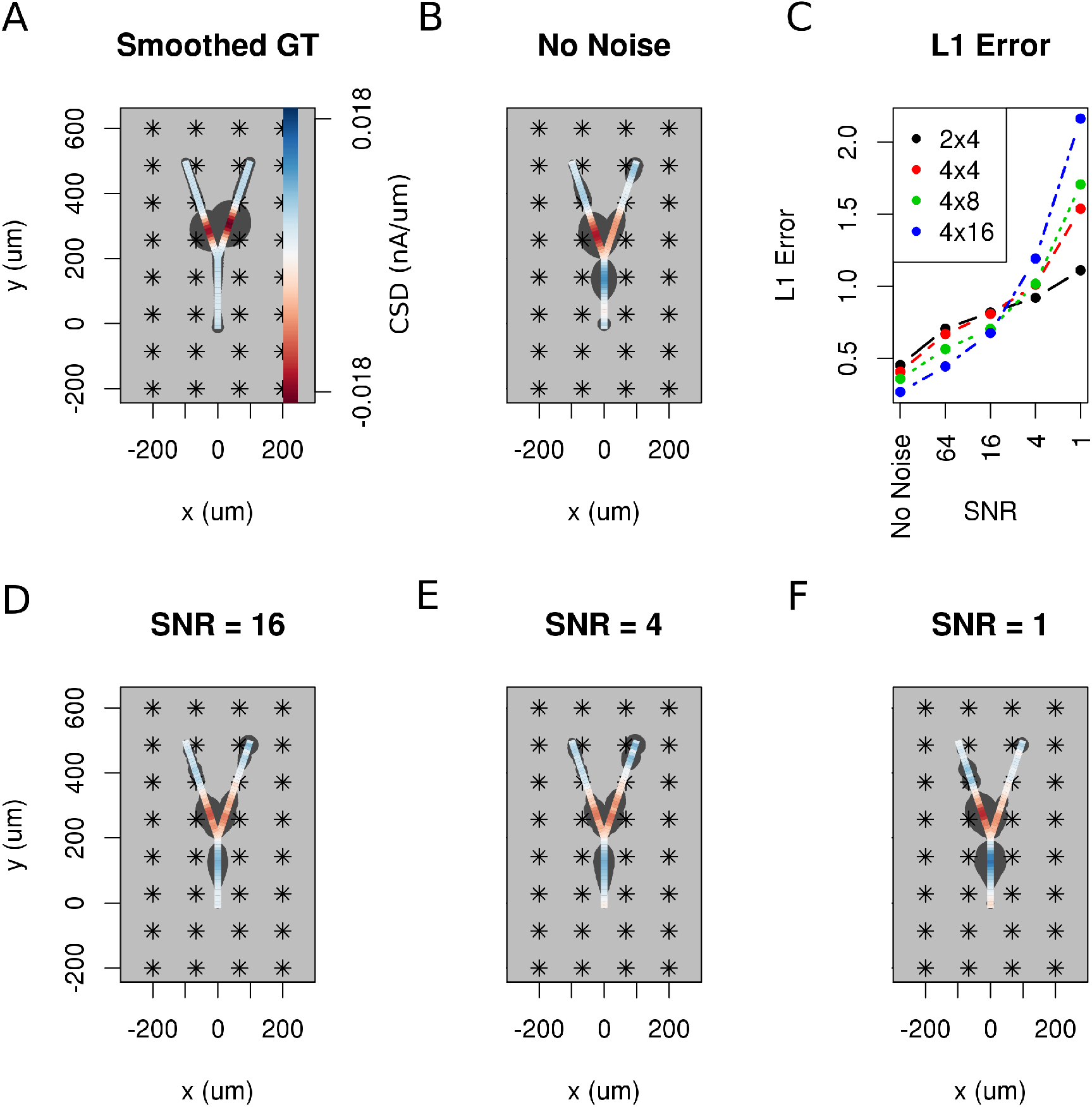
The effect of noise on the reconstruction. The corrupting influence of noise on the skCSD reconstruction is shown with the example of simultaneous excitation of both branches of the Y-shaped cell close to the branching point in case of the 4×8 electrodes setup. **A** Smoothed ground truth CSD shown on the branching morphology used. **B,D,E,F** skCSD reconstructions in cases of no added noise and signal-to-noise ratio equal to 16, 4, 1, respectively. Even the highest noise consider does not fully disrupt the reconstructed source distribution, although increasing the noise systematically degrades the result. This is shown in **C**, where the L1 error of the reconstruction was calculated for the full length of the simulations. This is consistent for different electrode setups which are marked with various colors. While the setups consisting of more electrodes perform better for low noise, the reconstruction seems to be more sensitive to noise in these cases. This might be a side effect of a specific definition of error.

### 3.5 Dependence of reconstruction on the number and arrangement of Recording Electrodes

Reconstruction of the distribution of the current sources along the morphology with skCSD, just like the reconstructions of smooth population distributions with kCSD, formally can be attempted from arbitrary set of recordings, even a single electrode. While we do not expect enlightening results at this extreme, it is natural to ask to what extent can we trust the reconstruction in a given case, which of the reconstructed features are real and which are artifacts of the method, and how to select optimal parameters of the method. We will discuss these issues in the final section. Here we wish to investigate how the number of electrodes, the density of the grid, and the area covered by the MEA, affect the results.

To answer these questions, we selected a snapshot of simulation of the model of the ganglion cell described in the Methods section, with the specific membrane current distribution shown in Fig. 9.A. In Fig. 9.B–H we show 7 different reconstructions assuming different experimental setups, with differing numbers of electrodes, covering different area.

**Figure 9.**
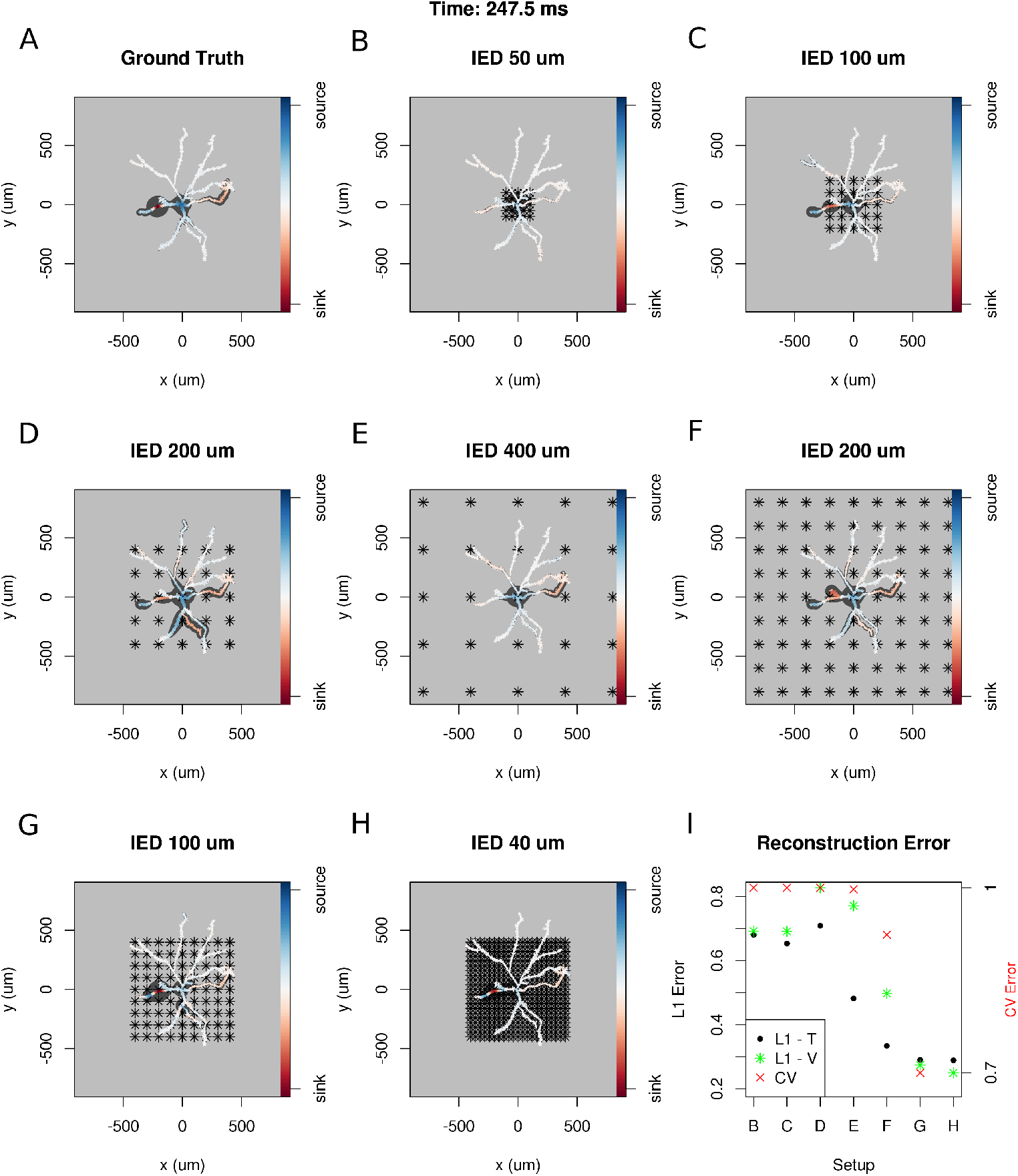
Dependence of skCSD reconstruction on multielectrode setup. Figures A–H show morphology used in the simulation together with the distribution of current sources in branching morphology representation taken at 247.5 ms of the simulation. Figures B–H show additionally the electrode setup assumed. **A**. Smoothed ground truth CSD. **B**. Reconstructed sources for a setup of 5x5 electrodes with 50 *μ*m interelectrode distance (IED) covering a small part of the cell morphology around the soma. **C**. Reconstructed sources for a setup of 5×5 electrodes with 100 *μ*m IED covering a substantial part of the dendritic tree, which improves the reconstruction of the synaptic input on the left. **D**. Reconstructed sources for 5×5 setup with 200 *μ*m IED setup; both sinks in the membrane currents are visible. **E**. Expanding the 5×5 electrode setup to 400 *μ*m IED leads to a small number of electrodes placed in the vicinity of the cell which leads to a poor reconstruction. **F**. Increasing the number of electrodes to 9×9 while keeping the coverage, which leads to 200 *μ*m IED, does not improve the reconstruction. **G**. Reducing IED in the previous example to 100 *μ*m, which reduces the coverage of the MEA to the whole cell (same area as in panel **D**) bringing majority of the electrodes close to one of the dendrites, leads to one of the most faithful reconstructions among the ones shown in this figure. **H**. Shows results for a matrix of 21×21 contacts with 40 *μ*m IED, covering the same area as in examples **D** and **G**. The results are very good but the improvement in reconstruction does not justify the use of so many contacts with so high density. **I**. Comparison of reconstruction errors for all the cases shown. Left axis: L1 error for the training (L1-T) and validation (L1-V) part. Right axis: crossvalidation error (CV). The L1-T error is marked with black points, L1-V error is represented by green stars. Generally, the L1-V errors are a bit higher than the L1-T errors but show a similar tendency. Also the CV errors, which are drawn with red crosses, show a similar tendency. The reconstructions in panels **B–H** are for parameters determined with the L1-T error.

In each case we selected the width of basis functions and the regularization parameter for the method by minimizing L1 error calculated for the first 1000 time steps of the simulation or cross-validation error (L1-T and CV in Fig. 9.I). To verify the quality of reconstruction we computed the L1 error between the ground truth and reconstruction for the remaining 5800 time steps of the simulation. It turns out that minimization of L1 error gave better results and L1-V in Fig. 9.I shows the results for this case.

Given that L1 error can only be used in simulations, where the ground truth is known, and yet it gives better parameters for the method to be applied to actual experimental data than the CV error, we propose the following. Given data necessary to apply skCSD method, thus the morphology, electrode positions, and recordings, one should assume different current sources distributions, for example, make a simulation of a cell model with the obtained morphology, make reconstructions for a range of parameters, and use L1 error for optimization. Then, perform the analysis of actual experimental data with thus obtained parameters. Performing the simulations and comparing the best reconstructions with the assumed ground truth has the further benefit of building intuition about which features of the real CSD survive in the reconstruction and which are distorted. This is another example of model-based data analysis which we believe becomes inevitable with the growing complexity of experimental paradigms, such as the one considered here.

The results obtained in this study are consistent with our expectations: the quality of reconstruction improves with the coverage of the morphology by the electrodes, with increasing density of probing, and with increasing number of probes (Fig. 9.I). Interestingly, it seems, that it is difficult to improve the reconstruction beyond certain level, in consequence, the setups with moderate densities (on the order of 200 μm IED) can easily compete with setups at the edge of current developments (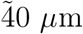 IED, [5]). We believe this is not a hard limit, that better results can be obtained here. This, however, requires further development of the methods.

### 3.6 Proof of Concept experiment: Spatial Current Source Distribution of Spike-triggered Averages

To examine the experimental feasibility of the skCSD method we analyzed data from a setup including simultaneous patch clamp electrode and linear probe with 14 working electrodes recording signals from a hippocampal pyramidal cell in vitro slice preparation (see Methods). As there is no ground truth data available in this case, the optimal width of the basis functions and the regularization parameter were selected using the L1 error and simulated data. To do this, we used the same simulation protocol as for the ganglion cell model. A snapshot of the reconstruction is shown in Fig. 10 at the moment of firing. A 10 ms long video of the spike triggered average is shown in the supplementary materials (S2 Video). At -0.05 ms one can observe a brief appearance of a sink (red) in the basal dendrites which can be a consequence of the activation of voltage sensitive channels in the axon hillock or the first axonal segment leading to the firing of the cell. Since the axon has not been traced in this case, the skCSD method is trying to reconstruct in the most meaningful way introducing the activity in the basal dendrite. This phenomenon is quickly replaced by a source (blue) in the basal dedrite with a sink at the soma and in the proximal part of the apical dendritic tree, with return current sinks at more distal dendrites. The extracellular potential on the second electrode reaches its minimum at 0.45 ms, which signals the peak of the spike. The deep red of the soma at this point signifies a strong sink, while the blue of the surrounding parts of the proximal apical and basal dendrites indicates the current sources set by the return currents. At 1.30 ms a source appears at the soma region, which indicates hyperpolarization. From an experimental setup consisting of only 14 electrodes on a linear probe a detailed distribution of current sources along a complex morphology cannot be expected, but the firing activity is well observable. This example demonstrates the experimental feasibility of the skCSD method and may help in planning further experiments using this method.

**Figure 10.**
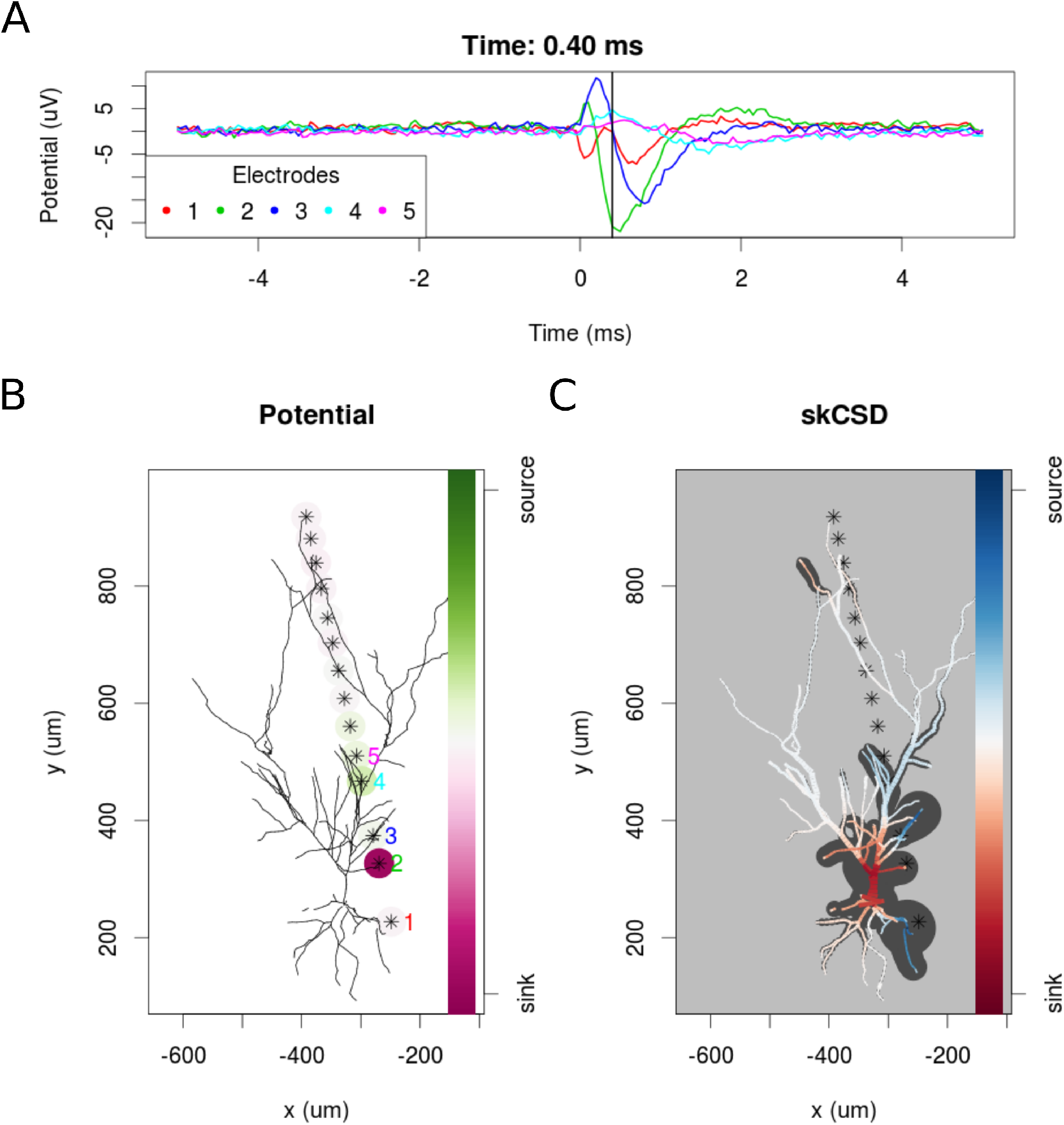
skCSD reconstruction of spike-triggered average for a hippocampal pyramidal cell. **A** Extracellular potentials measured with the 5 electrodes closest to the soma. The 0s marks the time of the membrane potential crossing the 0 mV threshold. The black vertical line marks the 0.40 ms time instant for which the extracellular potentials and skCSD reconstruction are shown. **B** Two-dimensional projection of the cell morphology and extracellular electrodes’ positions marked by stars, the five electrodes used in the top panel of the figure are labeled with matching colors. The amplitudes of the measured potentials are shown as color-coded circles around the electrodes. **C** The skCSD reconstruction on the branching morphology representation. This is a snapshot of the cell firing, the red color indicates the sinks close to the soma, the blue marks the current sources on the dendrites.

## 4 Discussion

### Summary

In this work we introduced a method to estimate the distribution of current sources (CSD) along the dendritic tree of a neuron given its known morphology and a set of simultaneous extracellular recordings of potential generated predominantly by this cell. First, assuming the ball-and-stick neuron model and a laminar probe parallel to the cell, we studied the basic viability of the method. We showed that introducing more electrodes to cover the same area leads to the increase of spatial resolution of the method allowing us to reconstruct higher Fourier modes of the CSD generating the measured potentials (Fig. 3). In a dynamic scenario of multiple synaptic inputs impinging on the cell, higher density of probes leads to higher reconstruction precision allowing us to distinguish individual inputs (Fig. 4). Testing the reconstruction against the known CSD (the ground truth) shows a clear transition between faithful and poor reconstruction when the electrode distribution becomes too sparse to capture the fine detail of the CSD profile to be reconstructed (Fig. 3.E). Also in neurons of more complex morphologies we studied, the Y-shape and the ganglion cell, as expected, the reconstructed CSD profiles became more detailed with the increase of electrode number on a fixed area (Fig. 6 and 9).

Using the Y-shaped morphology we showed that i) synaptic inputs activating different dendrites can be separated, Fig. 5; ii) skCSD provides meaningful information about the membrane CSD in cases, when interpolated LFP and standard, population CSD analysis, are not informative, Fig. 5; iii) the reconstruction is not sensitive to a specific selection of electrode placement, Fig. 6 and 8; and iv) even significant additive noise (SNR=1) is not prohibitive for the reconstruction, Fig. 8.

Biologically, the most relevant example we considered was a ganglion cell model which we studied with virtual multi-electrode arrays of different designs. The MEAs used differed with the inter-electrode distances for the simulated setups, as well as in the area they covered, ranging from an area close to the soma to roughly four times the size of the square covering the whole morphology. The best results where obtained when we used the electrodes from the region covering closely the cell (9.G and H); reduction of interelectrode distance from 100*μm* to 40*μm* was less spectacular than selecting the electrodes from the smallest square covering the convex hull span by the morphology. Our study, assuming realistic cell morphology of the ganglion cell and commercially available MEA designs, as well as realistic cell activity showed that it is feasible to reconstruct the distribution of the current sources in realistic, noisy situations.

The skCSD method performed adequately for the proof of concept experimental data, even if the nature of the setup allowed only the reconstruction of the general features of the spike-triggered average spatio-temporal current source density distribution patterns.

Historically, the idea of investigating the membrane currents of single cells was first proposed in [21], however, it used simplified, linear neuron morphologies. An important preprocessing step proposed there was separating the single neuron’s contributions to the extracellular potentials from the background activity. The novelty of the skCSD method proposed here is in its use of actual neuronal morphologies and in the underlying algorithmic solutions based on the kCSD method [26] devised for the study of populations of neurons.

### Experimental Recommendations

To attempt experimental application of skCSD we must have 1) an identified cell of known morphology, and 2) a set of simultaneous extracellular recordings of electric potential generated by this cell. Each aspect poses its challenges, some of which have been addressed here. Once we have the necessary data the natural question is how to select the parameters of the method in the specific context of a given setup, specific morphology, and recordings. Our investigations above give some indications: the electrodes selected for analysis should essentially uniformly cover the area span by the cell; the width of a basis source should be on the order of mean nearest neighbor interelectrode distance (for essentially uniformly distributed electrodes). We feel, however, that the proper approach is to actually investigate the effects of the different parameters through simulations. This is a natural place to apply the *model-based validation of data analysis* [39]. Our suggestion is to build a computational model of the cell. We believe that for the purpose of parameter selection assuming passive membrane in the dendrites should be sufficient, but of course, more realistic biophysical information may be included, especially if available. The model cell may be stimulated with synaptic input, with current injected, or even specific profiles of ground truth CSD may be placed along the cell. Then the extracellular potential must be computed at points where the actual electrodes are placed in the experiment. One can then investigate the effects of different parameter values on reconstruction and, for the analysis of actual experimental data, select those parameters minimizing prediction error on test data. The advantage of this procedure is two-fold. First, we end up with a selection of parameters adapted for the specific problem at hand. Secondly, we build intuition regarding the interpretation of the results for our specific cell and setup. This approach is the only way to address arbitrary electrode-cell configurations and to see how much information we can extract in a given case. Finally, we found that the best way to identify optimal parameters for reconstruction is by minimizing the L1 error between the reconstruction and the ground truth. Since we cannot have the ground truth in an experiment but we can assume it in the model-based validation, this is another argument for the model-based validation approach. Obviously, to efficiently apply this technique one must have appropriate tools. We plan to develop and open framework facilitating such studies, meanwhile, the code used for the present study will be made available upon request.

### Challenges of recording extracellular potential and obtaining morphology from the same cell

Although recording extracellular potential with a MEA, filling up a neuron with a dye, and reconstructing its morphology, are standard experimental techniques, using them simultaneously remains a challenge due to the size of the experimental devices which need to be arranged within a small volume. Cells in the vicinity of the MEA can be filled up individually by intracellular or juxtacellular electrodes, or with bulk dying. Individual recording and dying with a glass electrode provides not only the morphology, but also unambiguous spike times, gividing an opportunity to determine the extracellular potential fingerprint of the recorded cell on the MEA. Although these would be favorable data, intracellular recording less than 100*μm* from the MEA is extremely challenging. Experimental setups featuring the necessary equipment already exist [40], but as far as we know, haven’t been used in this way. On the other hand, bulk dying techniques result in more filled neurons, although the quality of the dying, and thus the quality of the 3D morphology reconstructions, is considerably lower in these cases. Although there are methods for estimation of the cell position relative to the MEA ([21], [22]), association of multiple optically labeled neurons with the recorded extracellular spike patterns is still unsolved.

### Challenges of separating the activity of a single neuron from background

We propose two ways to separate the activity of a neuron from the background. If we can sort the spikes elicited by the neuron of interest we can calculate the spike-triggered averages of the potentials reducing all uncorrelated contributions. Unfortunately, in live tissue, contributions from neighboring cells will have some correlations due to shared inputs. Separation of the contribution of the neuron of interest from the correlated background can be obtained in two ways. One is decomposition of the activity into meaningful components, for example, our results show that the high amplitude correlated oscillatory background of hippocampal theta activity can be extracted with independent component analysis, allowing the determination of cell-type specific time course of the synaptic input [41]. An alternative is combining skCSD with population kCSD analysis, i.e., inclusion of basis sources covering not just the cell of interest but also the space covering the whole population. This will be the subject of further study. A second way to obtain the contributions to the extracellular potential from a specific cell is by driving the cell with intracellular current injection of known pattern, for example, with an oscillatory drive as we discussed (Fig. 7), and by averaging over multiple periods (event-based triggering). Again, further study is needed to establish efficiency of such a procedure in experiment.

### Challenges of using novel MEAs

Handling data from high density MEAs with thousands of electrodes will require further studies, as the large numbers of small singular values of the kernel matrix may introduce numerical sensitivity to the reconstruction. Also, optimal selection of electrodes in case of programmable MEAs merits further investigations. We believe it is best to address such issues when actual experiments are attempted.

### Importance of this work

Traditional electrophysiology has focused on the electrical potential, which is relatively easy to access, from intracellular recordings, all kinds of patch clamp, juxtacellular, to extracellular and voltage sensitive dyes [42]. While the relation of the actual measurement to the voltage at a point may significantly differ, often this is a reasonable interpretation, if needed one may always consider more realistic models of measurement, for example, average over the contact surface for extracellular electrodes, etc [27, 43].

Already in the middle of XXth century, Walter H. Pitts had realized that with recordings on regular grids one can approximate the Poisson equation to estimate the distribution of current sources in the tissue, which he did [15]. His approach assumed recordings on a regular 3D grid, which was challenging to obtain for some 60 years [30]. However, with the work of Nicholson and Freeman [28] 1D CSD analysis became attractive, as summarized by Ulla Mitzdorf [16]. In 2012 we proposed how to overcome the restriction of regular grids with a kernel approach which both allows to use arbitrary distribution of contacts and corrects for noise [26]. All the previous work, however, always assumed the contributions to the extracellular potential coming from the whole tissue and smooth in the estimation region.

In the present work we show for the first time how one can use a collection of extracellular recordings in combination with a cell morphology to estimate the current sources located on the cell contributing to the recorded potential. Since it is now feasible experimentally to obtain the relevant data, we believe that the method proposed here may find its uses to constrain the biophysical properties of the neuron membrane, facilitate checking consistency of morphology reconstruction, as well as guide new discoveries by offering a more global picture of the distribution of the currents along the cell morphology, giving a coherent view of the global synaptic bombardment and return currents within a cell.

## 5 Acknowledgments

Research supported by grants from the Polish Ministry of Science and Higher Education 2948/7.PR/2013/2, the Hungarian National Research, Development and Innovation Fund NKFIH K 113147 and NN 118902, and the Hungarian National Brain Research Program KTIA NAP 13-1-2013-0001 and KTIA-13-NAP-A-IV/1,2,3,4,6. We are grateful for Emese Pálfi and László Négyessy for the opportunity of using their Neurolucida setup at the Department of Anatomy, Histology and Embryology, Semmelweis University.

## Supporting Information

### S1 Video

#### skCSD reconstruction of current source density distribution on the ganglion cell

The video shows the skCSD reconstruction for the retinal ganglion cell model driven with oscillatory current (Section 3.3) for the whole duration of simulation. Figure 7 shows a snapshot taken at *t* = 495.25 ms from the simulation onset. As described in Section 3.3, during the first 400 ms of simulation, apart from somatic drive, 100 excitatory synaptic inputs were randomly distributed along the dendrites. For reconstruction, 128 virtual electrodes were selected from the 936 arranged in a hexagonal grid of 17.5 *μm* interelectrode distance to record the extracellular potentials. Panel A presents the somatic membrane potential during the simulation. The red line marks the time instant for which the remaining plots were made. The colormap on Panel B shows the extracellular potential interpolated between the simulated measurements computed at the electrodes, which are marked with asterisks. The regular CSD is shown on Panel C, while the spatially smoothed ground truth membrane current is presented on Panel D. Panel E shows the skCSD reconstruction of current source density along the cell morphology from the selected measurements. The dark gray shapes are guides for the eye and are sums of circles placed along the morphology with radius proportional to the amplitude of the sources at the center of the circle.

The movie is available at https://www.dropbox.eom/s/8ea0q9mjhgk2s0x/CSDSmoothedmp4?dl=0.

### S2 Video

#### Spike triggered average of pyramidal cell in vitro

The video shows the recorded potentials and skCSD reconstruction for a 10 ms time window centered around the spike as described in Section 2.7. The top panel presents the spike triggered averages of the potentials during 5 seconds before and after the spike recorded at five electrodes closest to the soma. The lower left panel shows the morphology of the cell, electrode positions, and the recorded potentials. The electrodes are marked by stars and the amplitude of the recorded potential is shown as color-coded circles around the electrodes. The snapshot is taken at the time given in the figure title and indicated by the black vertical line in the top panel. The reconstructed skCSD distribution at the same moment is shown in the lower right panel.

At −0.05 ms a sink appears at the basal dendrites. This can be a consequence of the activation of voltage sensitive channels in the axon hillock or the first axonal segment leading to the firing of the cell. Since there were no electrodes close to the axon initial segment, the skCSD method did not resolve it and reconstruct the source by introducing the activity into the basal dendrite. This phenomenon is quickly replaced by a source (blue) in the basal dedrite with a sink at the soma and in the proximal part of the apical dendritic tree, with return current sinks at more distal dendrites. The extracellular potential on the second electrode reaches its minimum at 0.45 ms, which signals the peak of the spike. The deep red of the soma at this point signifies a strong sink, while the blue of the surrounding parts of the proximal apical and basal dendrites indicates the current sources set by the return currents. At 1.30 ms a source appears at the soma region, which indicates hyperpolarization.

The movie is available at https://www.dropbox.com/s/50tgai6oiz172zm/PyramidalSTAmp4?dl=0.

## References

1. Neher E, Sakmann B. Single-channel currents recorded from membrane of denervated frog muscle fibres. Nature. 1976 Apr;260(5554):799–802.

2. Buzsaki G, Anastassiou C, Koch C. The origin of extracellular fields and currents–EEG, ECoG, LFP and spikes. Nature Reviews Neuroscience. 2012;13:407–420.

3. Einevoll GT, Kayser C, Logothetis NK, Panzeri S. Modelling and analysis of local field potentials for studying the function of cortical circuits. Nature Reviews Neuroscience. 2013;14:770–785.

4. Buzsáki G. Large-scale recording of neuronal ensembles. Nat Neurosci. 2004 May;7(5):446–451. Available from: http://dx.doi.org/10.1038/nn1233.

5. Berdondini L, van der Wal PD, Guenat O, de Rooij NF, Koudelka-Hep M, Seitz P, et al. High-density electrode array for imaging in vitro electrophysiological activity. Biosens Bioelectron. 2005 Jul;21(1):167–174. Available from: http://dx.doi.org/10.1016/j.bios.2004.08.011.

6. Obien MEJ, Deligkaris K, Bullmann T, Bakkum DJ, Frey U. Revealing neuronal function through microelectrode array recordings. Front Neurosci. 2014;8:423. Available from: http://dx.doi.org/10.3389/fnins.2014.00423.

7. Rey HG, Pedreira C, Quiroga RQ. Past, present and future of spike sorting techniques. Brain research bulletin. 2015;119:106–117.

8. Hottowy P, Skoczeń A, Gunning DE, Kachiguine S, Mathieson K, Sher A, et al. Properties and application of a multichannel integrated circuit for low-artifact, patterned electrical stimulation of neural tissue. Journal of neural engineering. 2012;9(6):066005.

9. Chichilnisky E. A simple white noise analysis of neuronal light responses. Network: Computation in Neural Systems. 2001;12(2):199–213.

10. Ferrea E, Maccione A, Medrihan L, Nieus T, Ghezzi D, Baldelli P, et al. Large-scale, high-resolution electrophysiological imaging of field potentials in brain slices with microelectronic multielectrode arrays. Front Neural Circuits. 2012;6:80. Available from: http://dx.doi.org/10.3389/fncir.2012.00080.

11. Bakkum DJ, Frey U, Radivojevic M, Russell TL, Müller J, Fiscella M, et al. Tracking axonal action potential propagation on a high-density microelectrode array across hundreds of sites. Nat Commun. 2013;4:2181. Available from: http://dx.doi.org/10.1038/ncomms3181.

12. Lewandowska MK, Radivojević M, Jäckel D, Müller J, Hierlemann AR. Cortical Axons, Isolated in Channels, Display Activity-Dependent Signal Modulation as a Result of Targeted Stimulation. Front Neurosci. 2016;10:83. Available from: http://dx.doi.org/10.3389/fnins.2016.00083.

13. Jäckel D, Bakkum DJ, Russell TL, Müller J, Radivojevic M, Frey U, et al. Combination of High-density Microelectrode Array and Patch Clamp Recordings to Enable Studies of Multisynaptic Integration. Scientific reports. 2017 Apr;7:978.

14. Muthmann JO, Amin H, Sernagor E, Maccione A, Panas D, Berdondini L, et al. Spike Detection for Large Neural Populations Using High Density Multielectrode Arrays. Front Neuroinform. 2015;9:28. Available from: http://dx.doi.org/10.3389/fninf.2015.00028.

15. Pitts WH. Investigations on synaptic transmission. In: Cybernetics, Trans. 9th Conf. Josiah Macy Foundation H. von Foerster. New York; 1952. p. 159–166.

16. Mitzdorf U. Current source-density method and application in cat cerebral cortex: investigation of evoked potentials and EEG phenomena. Physiological Reviews. 1985;65:37–100.

17. Wójcik DK. Current Source Density (CSD) Analysis. In: Jaeger D, Jung R, editors. Encyclopedia of Computational Neuroscience. Springer New York; 2015. p. 915–922. Available from: http://dx.doi.org/10.1007/978-1-4614-6675-8_544.

18. Einevoll GT, Lindén H, Tetzlaff T, Łęski S, Pettersen KH. Local field potential: biophysical origin and analysis. In: Quiroga RQ, Panzeri S, editors. Principles of Neural Coding. CRC Press; 2013. p. 37–59.

19. Głąbska H, Potworowski J, Łęski S, Wójcik DK. Independent components of neural activity carry information on individual populations. PLoS One. 2014;9(8):e105071. Available from: http://dx.doi.org/10.1371/journal.pone.0105071.

20. Głąbska HT, Norheim E, Devor A, Dale AM, Einevoll GT, Wójcik DK. Generalized Laminar Population Analysis (gLPA) for Interpretation of Multielectrode Data from Cortex. Front Neuroinform. 2016;10:1. Available from: http://dx.doi.org/10.3389/fninf.2016.00001.

21. Somogyvári Z, Zalányi L, Ulbert I, Erdi P. Model-based source localization of extracellular action potentials. J Neurosci Methods. 2005 Sep;147(2):126–137. Available from: http://dx.doi.org/10.1016/j.jneumeth.2005.04.002.

22. Somogyvári Z, Cserpán D, Ulbert I, Erdi P. Localization of single-cell current sources based on extracellular potential patterns: the spike CSD method. Eur J Neurosci. 2012 Nov;36(10):3299–3313. Available from: http://dx.doi.org/10.1111/j.1460-9568.2012.08249.x.

23. Mechler F, Victor JD, Ohiorhenuan I, Schmid AM, Hu Q. Three-dimensional localization of neurons in cortical tetrode recordings. J Neurophysiol. 2011 Aug;106(2):828–848. Available from: http://dx.doi.org/10.1152/jn.00515.2010.

24. Mechler F, Victor JD. Dipole characterization of single neurons from their extracellular action potentials. J Comput Neurosci. 2012 Feb;32(1):73–100. Available from: http://dx.doi.org/10.1007/s10827-011-0341-0.

25. Frey U, Egert U, Heer F, Hafizovic S, Hierlemann A. Microelectronic system for high-resolution mapping of extracellular electric fields applied to brain slices. Biosens Bioelectron. 2009 Mar;24(7):2191–2198. Available from: http://dx.doi.org/10.1016/j.bios.2008.11.028.

26. Potworowski J, Jakuczun W, Łęski S, Wójcik D. Kernel current source density method. Neural Comput. 2012 Feb;24(2):541–575. Available from: http://dx.doi.org/10.1162/NECO_a_00236.

27. Ness TV, Chintaluri C, Potworowski J, Łęski S, Głąbska H, Wójcik DK, et al. Modelling and analysis of electrical potentials recorded in microelectrode arrays (MEAs). Neuroinformatics. 2015;13(4):403–426.

28. Nicholson C, Freeman JA. Theory of current source-density analysis and determination of conductivity tensor for anuran cerebellum. J Neurophysiol. 1975 Mar;38(2):356–368.

29. Pettersen KH, Devor A, Ulbert I, Dale AM, Einevoll GT. Current-source density estimation based on inversion of electrostatic forward solution: effects of finite extent of neuronal activity and conductivity discontinuities. J Neurosci Methods. 2006 Jun;154(1–2):116–133. Available from: http://dx.doi.org/10.1016/j.jneumeth.2005.12.005.

30. Łęski S, Wójcik DK, Tereszczuk J, Świejkowski DA, Kublik E, Wróbel A. Inverse current-source density method in 3D: reconstruction fidelity, boundary effects, and influence of distant sources. Neuroinformatics. 2007;5(4):207–222. Available from: http://dx.doi.org/10.1007/s12021-007-9000-z.

31. Łęski S, Pettersen KH, Tunstall B, Einevoll GT, Gigg J, Wójcik DK. Inverse current source density method in two dimensions: inferring neural activation from multielectrode recordings. Neuroinformatics. 2011 Dec;9(4):401–425. Available from: http://dx.doi.org/10.1007/s12021-011-9111-4.

32. Kwan MK. Graphic programming using odd or even points. Chinese Math. 1962;1(273–277):110.

33. Lindén H, Hagen E, Lęski S, Norheim ES, Pettersen KH, Einevoll GT. LFPy: a tool for biophysical simulation of extracellular potentials generated by detailed model neurons. Front Neuroinform. 2013;7:41. Available from: http://dx.doi.org/10.3389/fninf.2013.00041.

34. Cannon RC, Turner DA, Pyapali GK, Wheal HV. An on-line archive of reconstructed hippocampal neurons. Journal of Neuroscience Methods. 1998;84(1–2):49–54. Available from: http://www.sciencedirect.com/science/article/pii/S0165027098000910.

35. Kong JH, Fish DR, Rockhill RL, Masland RH. Diversity of ganglion cells in the mouse retina: unsupervised morphological classification and its limits. J Comp Neurol. 2005 Aug;489(3):293–310. Available from: http://dx.doi.org/10.1002/cne.20631.

36. Ascoli GA. Mobilizing the base of neuroscience data: the case of neuronal morphologies. Nat Rev Neurosci. 2006 Apr;7(4):318–324. Available from: http://dx.doi.org/10.1038/nrn1885.

37. Hansen P. Discrete Inverse Problems. Society for Industrial and Applied Mathematics; 2010. Available from: http://epubs.siam.org/doi/abs/10.1137/1.9780898718836.

38. Kerekes BP, Tóth K, Kaszás A, Chiovini B, Szadai Z, Szalay G, et al. Combined two-photon imaging, electrophysiological, and anatomical investigation of the human neocortex in vitro. Neurophotonics. 2014;1(1):11013. Available from: http://dx.doi.org/10.1117/1.NPh.1.1.011013.

39. Denker M, Einevoll G, Franke F, Grün S, Hagen E, Kerr J, et al. Report from the 1st INCF Workshop on Validation of Analysis Methods. INCF; 2014.

40. Neto JP, Lopes G, Frazão J, Nogueira J, Lacerda P, Baião P, et al. Validating silicon polytrodes with paired juxtacellular recordings: method and dataset. Journal of Neurophysiology. 2016;116(2):892–903. Available from: http://jn.physiology.org/content/116/2/892.

41. Somogyvári Z, Benkő Z, Jálics JZ, Roux L, Antal B. Determination of spatio-temporal input current patterns of single hippocampal neurons based on extracellular potential measurements. Program No. 267.02.. Society for Neuroscience, 2015; 2015.

42. Covey E, Carter M. Basic Electrophysiological Methods. Oxford University Press, Incorporated; 2015. Available from: https://books.google.pl/books?id=-udcBgAAQBAJ.

43. Moulin C, Glière A, Barbier D, Joucla S, Yvert B, Mailley P, et al. A new 3-D finite-element model based on thin-film approximation for microelectrode array recording of extracellular action potential. IEEE transactions on bio-medical engineering. 2008 feb;55(2 Pt 1):683–92. Available from: http://www.ncbi.nlm.nih.gov/pubmed/18270005.

